# Protein features for assembly of the RNA editing helicase 2 subcomplex (REH2C) in Trypanosome holo-editosomes

**DOI:** 10.1101/524702

**Authors:** Vikas Kumar, Pawan K. Doharey, Shelly Gulati, Joshua Meehan, Mary G. Martinez, Karrisa Hughes, Blaine H.M. Mooers, Jorge Cruz-Reyes

## Abstract

Uridylate insertion/deletion RNA editing in *Trypanosoma brucei* is a complex system that is not found in humans, so there is interest in targeting this system for drug development. This system uses hundreds of small non-coding guide RNAs (gRNAs) to modify the mitochondrial mRNA transcriptome. This process occurs in holo-editosomes that assemble several macromolecular *trans* factors around mRNA including the RNA-free RNA editing core complex (RECC) and auxiliary ribonucleoprotein (RNP) complexes. Yet, the regulatory mechanisms of editing remain obscure. The enzymatic accessory RNP complex, termed the REH2C, includes mRNA substrates and products, the multi-domain 240 kDa RNA Editing Helicase 2 (REH2) and an intriguing 8-zinc finger protein termed REH2-Associated Factor 1 (^H2^F1). Both of these proteins are essential in editing. REH2 is a member of the DExH/RHA subfamily of RNA helicases with a conserved C-terminus that includes a regulatory OB-fold domain. In trypanosomes, ^H2^F1 recruits REH2 to the editing apparatus, and ^H2^F1 downregulation causes REH2 fragmentation. Our systematic mutagenesis dissected determinants in REH2 and ^H2^F1 for the assembly of REH2C, the stability of REH2, and the RNA-mediated association of REH2C with other editing *trans* factors. We identified functional OB-fold amino acids in eukaryotic DExH/RHA helicases that are conserved in REH2 and that impact the assembly and interactions of REH2C. ^H2^F1 upregulation stabilized REH2 *in vivo*. Mutation of the core cysteines or basic amino acids in individual zinc fingers affected the stabilizing property of ^H2^F1 but not its interactions with other examined editing components. This result suggests that most, if not all, fingers may contribute to REH2 stabilization. Finally, a recombinant REH2 (240 kDa) established that the full-length protein is a *bona fide* RNA helicase with ATP-dependent unwinding activity. REH2 is the only DExH/RHA-type helicase in kinetoplastid holo-editosomes.

## Introduction

Kinetoplastid protozoa including *Trypanosoma brucei* are early-branching eukaryotes that require extensive insertion and deletion of uridylates in most mitochondrial mRNAs (reviewed in 1). This RNA editing is protein-catalyzed and directed by small guide RNAs (gRNAs) with complementarity to fully-edited mRNA. Most editing progresses in blocks that typically overlap and each block is directed by a single gRNA (2). The editing machinery includes multiple subcomplexes: the ∼20S catalytic RNA editing core complex (RECC, also called the 20S editosome) and auxiliary editing RNPs. The ∼15S REH2-associated complex (REH2C) characterized here includes RNA editing helicase 2 (REH2), REH2-associated protein factor 1 (^H2^F1), ^H2^F2, and mRNA substrates and products (3, 4). However, only REH2 and ^H2^F1 participate in editing. REH2C interacts with a larger accessory editing RNP complex termed the RNA editing substrate binding complex (RESC), which also contains mRNA (5, 6). The presence of mRNA in RESC was confirmed by others (7, 8).

RESC contains two modules that are intimately associated with each other: the gRNA-binding complex (GRBC) typified by the GAP1/GAP2 (alias GRBC2/GRBC1) heterotetramer that binds and stabilizes gRNA, and the more loosely defined RNA editing mediator complex (REMC) (6). REMC is typified by RGG2 and promotes editing progression (8–10). Other mRNA *trans* factors include a subcomplex containing MRB6070 and MRB1590, and RNA helicase REH1 that associate with holo-editosomes via stable or transient RNA contacts, respectively (11–13). The finding that the editing RNPs carry mRNA and gRNA implied that mRNA-gRNA hybrids form on multi-RNP platforms, and that transient addition of the RECC enzyme to the substrate-loaded platforms completes the assembly of the holo-editosome (3–8).

The editing apparatus is further complicated by the occurrence of variants of GRBC, REMC and RECC that may be regulatory (reviewed in 1) but the precise control mechanisms in editing remain unclear. However, the helicase REH2C subcomplex may play an important regulatory role. REH2 is the largest characterized member of the DExH/RNA helicase A (RHA) subfamily of proteins (14). These proteins are monomeric and contain a conserved C-terminal domain cluster with a characteristic auxiliary oligonucleotide-binding (OB fold) domain (3, 14-16). The OB fold in some RNA helicases is known to bind protein regulators or mediate RNA-dependent activation of these enzymes (17). DExH/RHA proteins have a N-terminus with a variable domain organization of less clear function. The N-terminal half of the REH2 carries two predicted double-stranded RNA binding domains: dsRBD1 and dsRBD2 (14). Other characterized DExH/RHA-type proteins that carry two N-terminal dsRBDs are the RNA helicase RHA (aka DHX9) in vertebrates and its orthologous RNA helicase MLE (*maleless*) in flies (18). Recombinant versions of the REH2 C-terminal half and full-size octa-zinc finger ^H2^F1 formed a stable complex *in vitro* (4). However, ^H2^F1 is an unusual binding partner of DExH/RHA proteins that typically associate with G-patch proteins (17). A genetic knockdown of ^H2^F1 prevented RNA-mediated association of REH2 with other editing components and caused fragmentation of REH2 *in vivo*. Those results indicated that ^H2^F1 is an adaptor protein and may stabilize REH2 (4).

The current study identified features in REH2 and ^H2^F1 that affect the binding of these proteins with each other, the stability of REH2, and RNA-mediated stable or transient association of the REH2C subcomplex with other editing components *in vivo*. About 20 different protein variants were examined *in vivo*. Comparisons of the REH2 with other eukaryotic DExH/RHA helicases identified conserved OB fold residues that affect the assembly and interactions of REH2C and are also important in distant processes, namely mRNA splicing in yeast, and the assembly of the dosage compensation complex in fly. ^H2^F1 upregulation stabilized the large REH2 polypeptide *in vivo*. However, mutation of individual zinc fingers compromised the stabilizing property of ^H2^F1 but not its association with REH2 or other examined components of the editing apparatus. Most if not all fingers may contribute to REH2 stabilization. Finally, we used a recombinant protein construct to establish that the isolated full-length REH2 (240 kDa) is a *bona fide* RNA helicase enzyme with ATP-dependent unwinding activity. REH2C is most likely an enzymatic editing RNP in kinetoplastid holo-editosomes.

## Results

Previous studies revealed that the native REH2C exhibits RNA-mediated interactions with at least two variants of RESC (3–5). These variants contain gRNA and comparable levels of GAP1 and RGG2, that typify GRBC and REMC, respectively. However, these variants differ minimally in their relative content of the canonical MRB3010 in GRBC. Additional variations in protein composition have been reported in purifications of RESC components by different labs (reviewed in 1). Interestingly, native and genetically-induced changes in MRB3010 stoichiometry may be common. While the precise reasons for these changes are unclear, we reasoned that they reflect relevant dynamic changes in the assembly and function of the editing apparatus. In this study, we examined protein features that affect the formation of REH2C, its stable association with RESC components, and its transient contacts with the RECC enzyme. Because the examined helicase-associated RESC variant exhibits substoichiometric MRB3010, we refer to its variant GRBC module as GRBC* (4). Using antibodies made available to us, we have detected other proteins in pulldowns of REH2 that may be part of this RESC variant including both GAP1/GAP2 paralogs and MRB8170. The REH2 pulldowns also contain MRB6070 that was found in a MRB6070/MRB1590 subcomplex that associates via RNA with RESC (11)(S1 Fig.). However, further analysis of relative composition between variants is needed.

### Analysis of REH2 mutants *in vivo*

We analyzed the importance of specific features of REH2, both in the formation of REH2C and in the association of this subcomplex with other components in holo-editosomes (constructs listed in Fig. 1A). To this end, we expressed a tetracycline (Tet)-inducible REH2 wild-type (WT) construct with a Tandem Affinity Purification (TAP) tag (3) and examined its co-purification with endogenous ^H2^F1 and GAP1 proteins, of REH2C and GRBC, respectively. We found that ectopic REH2 was able to reconstitute interactions of the native REH2 protein in trypanosomes (Fig. 1B-C, lanes 1-2, respectively). With this system in hand, we tested the assembly of REH2 variants bearing truncations or point mutations in an approach that has been used in other eukaryotic DExH/RHA helicases (16, 19). All REH2 mutants examined were directly compared and normalized to a WT construct used as control. Neither overexpression or depletion of REH2 affects the steady-state level of its binding partner ^H2^F1 or the examined GRBC proteins (3, 4).

**Fig. 1.**
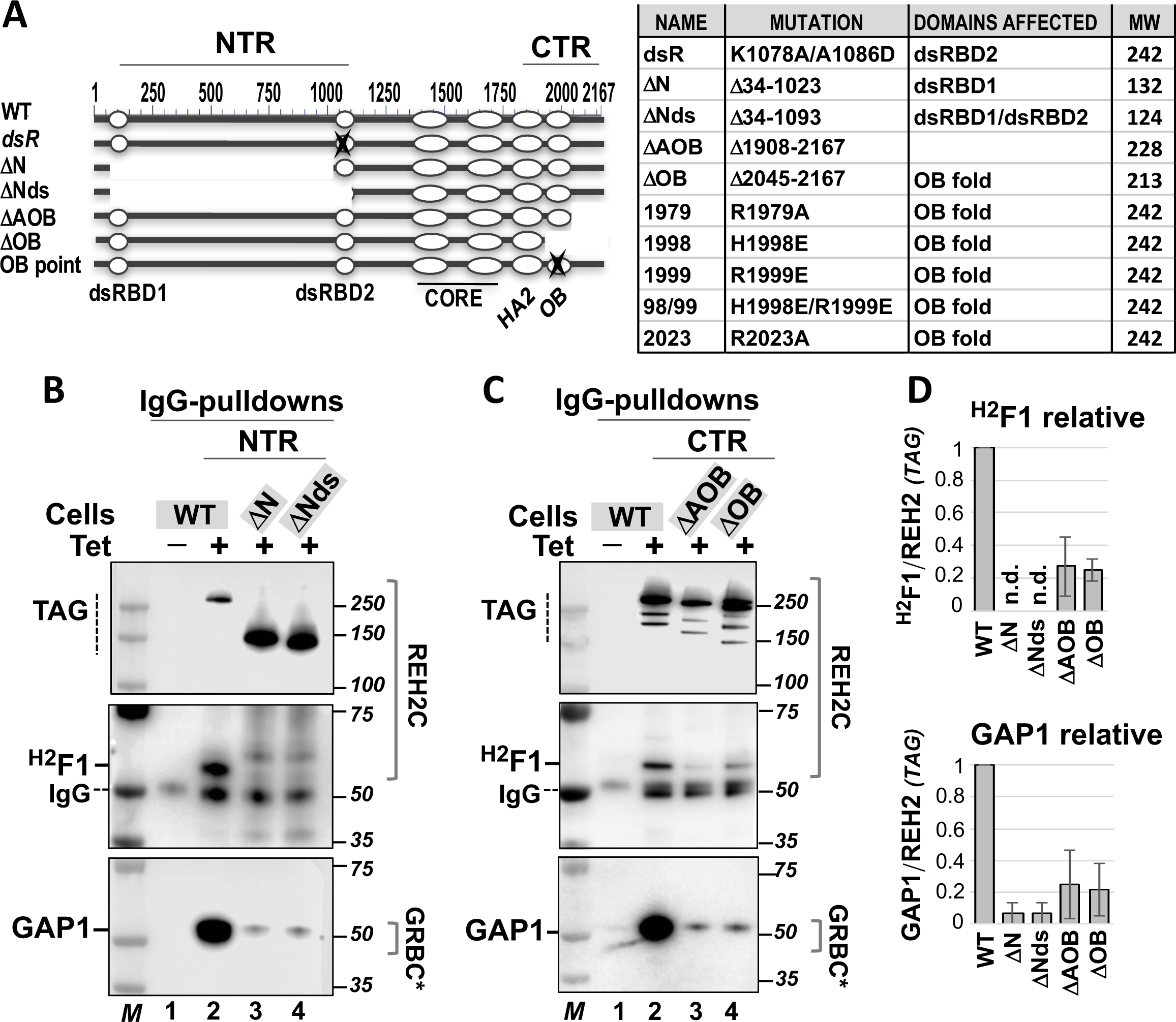
*In vivo* analyses of REH2 variants. **(A)** Scheme of REH2 WT (2167 residues) and variants with the examined truncations or point mutations in the N- and C-terminal regions (NTR and CTR). The mutations, affected domains, and predicted molecular weight of the constructs are listed. Identified domains: dsRBD1 and dsRBD2 (double-stranded RNA binding domains 1 and 2), CORE (catalytic core RecA1 and RecA2 domains), HA2 (helicase associated helical bundle domain), and OB (OB fold-like domain). **(B-C)** Western blots of IgG pulldowns from extracts to examine the association of tagged REH2 WT or deletion mutants with endogenous ^H2^F1 and GAP1 (in GRBC*). An uninduced (-Tet) control lane is included. The data are representative of at least three biological replicates for each mutant. A few non-specific species cross-react with the antibodies in some assays. The position of IgG is marked. Sizing markers (M) are in kDa. **(D)** Charts of the relative level of ^H2^F1 and GAP1 in the pulldowns. Representative assays from panels A-B and other pulldowns in this study were used to plot +/-1 SD, *n*=3 for each mutant. The ^H2^F1/REH2 and GAP1/REH2 ratios in each REH2 mutant were normalized to the ratios in REH2 WT (= 1, in the plot). Assays with signals that were too low were not determined (n.d.). REH2 was often fragmented, so only the full-length polypeptide was scored. Some deletion mutants were present at higher level than tagged REH2 WT in the induced extracts.

#### N- and C-truncations

We examined N- and C-terminal regions (NTR and CTR) flanking the catalytic and helicase associated domains RecA1/RecA2 and HA2, respectively, in REH2. We removed sequences in the unique NTR (>1000 residues) in the REH2 variants ΔN and ΔNds. Next, we removed the CTR after the OB fold (123 residues) or from the OB fold (260 residues) in the ΔAOB and ΔOB variants, respectively. REH2 and its orthologs in kinetoplastids carry the longest N terminus in the known DExH/RHA-type helicases (∼1300 residues in REH2 from *T. brucei*) (14). Direct comparisons with the REH2 WT construct showed that the examined N- and C-resections reduced the association of the shortened REH2 polypeptides with ^H2^F1 and GAP1 proteins (Figs. 1B-D). This suggested that both termini of REH2 may contribute to the normal assembly of REH2C and its RNA-mediated interactions with other editing components *in vivo*. The dsRBDs, the OB fold, and undefined features in the N and C termini may be involved. It is also conceivable that the examined sequence resections may impact the global conformation of REH2, thereby indirectly affecting its protein interactions. In that case, the normal REH2 interactions in the editing apparatus may be sensitive to overall changes in the integrity or conformation of REH2. A comparison of REH2 ΔN and ΔNds in whole-cell and enriched-mitochondrial extracts indicated that these large truncations do not prevent mitochondrial import (S2 Fig). Because the CTR includes a potentially regulatory OB fold in DExH/RHA-type RNA helicases (16), we focused on this domain in the subsequent studies of REH2.

#### Point mutations

Our structural searches of the OB fold-like domain in REH2 using Phyre2 (20) found the best agreements of the REH2 OB fold with structures of yeast ADP-bound Prp43p helicase and *Drosophila* RNA helicase MLE in a complex with RNA and ADP-Alf4 (4, 16, 18, 21). Prp43p is a bi-functional helicase in mRNA splicing and ribosome biogenesis (15, 22). The helicase MLE remodels roX RNA substrates to promote assembly of the dosage compensation complex in *Drosophila* (18). We examined specific residues in the predicted OB fold of REH2 that seem to be conserved in the DExH/RHA helicases mentioned above (Fig. 2 A-D). OB point mutations were compared to two controls: REH2 WT and a mutation in dsRBD2 (dsR) that reduces the association of REH2 with GAP1, gRNA and mRNA (Fig. 2A, lanes 1 and 2, respectively) (3). The β1−β2 and β4−β5 loops in the OB fold may include determinants in nucleic acid recognition (Fig. 2D) (16, 18). In helicase Prp43p, mutation of K704 (β4−β5 loop), which is exposed to the solvent in the Prp43p structure, decreased the RNA affinity and ATPase activity of the helicase *in vitro* (16). Sequence alignments and a homology model derived from the structure of Prp43p matched K704 in Prp43p with R2023 in the predicted β4−β5 loop in REH2 (S3 Fig)(4). We also examined R1979 in REH2 that is not conserved in Prp43p and may serve as an internal control. Pulldowns of the R1979A and R2023A mutants suggest that neither mutation alters the association of REH2 with ^H2^F1 (Figs. 2A-C). R1979A did not appear to affect the REH2 association with GAP1. However, R2023A caused a moderate decrease in the association with GAP1, implying that at least R2023A could be disruptive in trypanosomes.

**Fig. 2.**
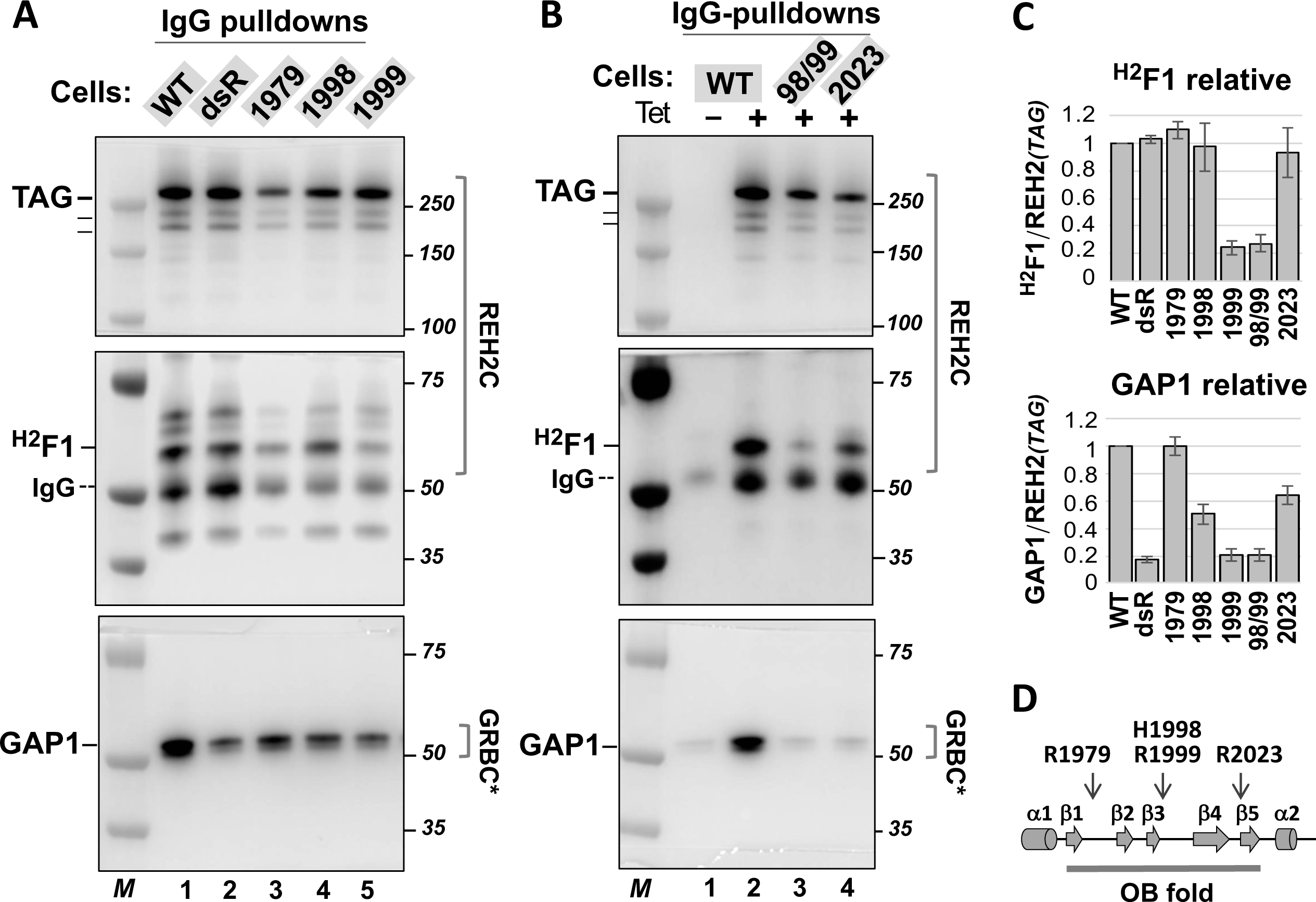
*In vivo* analyses of REH2 variants. Western blots of IgG pulldowns as in Fig. 1 to examine the association of endogenous ^H2^F1 and GAP1 (in GRBC*) with tagged REH2 WT or variants with point mutations at: **(A)** the OB fold domain including R1979A (1979), H1998E (1998) and R1999E (1999). The previously characterized dsR mutant with an inactivating double mutation in the dsRBD2 was included for comparison (3). A few non-specific species crossreact with the antibodies in some assays; and **(B)** the OB fold domain including double 1998/1999 (98/99) or single R2023A (2023) mutations. **(C)** Charts of the relative levels of ^H2^F1 and GAP1 in the pulldowns. Assays from panels A-B and biological replicate pulldowns in this study were used to plot +/-1 SD, *n*=2 or *n*=3 for each mutant. The ^H2^F1/REH2 and GAP1/REH2 ratios in each REH2 mutant were normalized to the ratios in REH2 WT (= 1, in the plot). **(D)** Secondary structure prediction of the OB fold with *α*-helix (cylinders) and β-strand (arrows) elements, and the position of point mutations examined indicated by arrows.

In helicase MLE, several residues in the OB fold bind to specific uridylates in U-rich sequences in roX transcripts. Mutation of some of these residues impaired RNA binding by MLE *in vitro* or the proper localization of MLE in chromosomes (18). Sequence alignments and homology models indicated that the residues H1032 and K1033 in MLE likely represent residues H1998 and R1999 in the predicted β3−β4 loop in the OB fold in REH2 (Fig. 2D; S4 Fig)(21). The H1998E mutant appeared to retain a normal association with ^H2^F1 but exhibited a moderate decrease in association with GAP1. In contrast, R1999E either alone or together with H1998E (98/99) exhibited a more robust decrease (over 60%) in association with both ^H2^F1 and GAP1 (Fig. 2A-C). This indicated that H1998 and R1999 contribute to the assembly of the REH2 helicase with other editing components. However, R1999 has a higher impact on such interactions than the adjoining H1998. As expected, the dsR mutant used as a control exhibited a strong decrease in association with GAP1. However, the dsR mutation did not seem to affect the association of REH2 with ^H2^F1 (Figs. 2A, 2C). Pulldowns of the mutants were normalized to the REH2 WT control as in Fig. 1.

Table 1 summarizes observed effects of the examined mutations in REH2 on its association with ^H2^F1 and GAP1. While large protein truncations in REH2 may alter the global conformation of the polypeptide, relevant point mutations in REH2 are more likely to cause specific functional effects. Our systematic mutagenesis of REH2 suggests an intricate interaction network including features across both N- and C-terminal regions of REH2 that affect the assembly of REH2C, and the association of this subcomplex with GAP1-associated RNPs *in vivo*. Remarkably, these observations also indicate that equivalent residues in the auxiliary conserved OB fold domain in DExH/RHA helicases function in disparate RNA processes in the taxonomically distant species of trypanosomes, yeast, and fly.

**Table 1.**
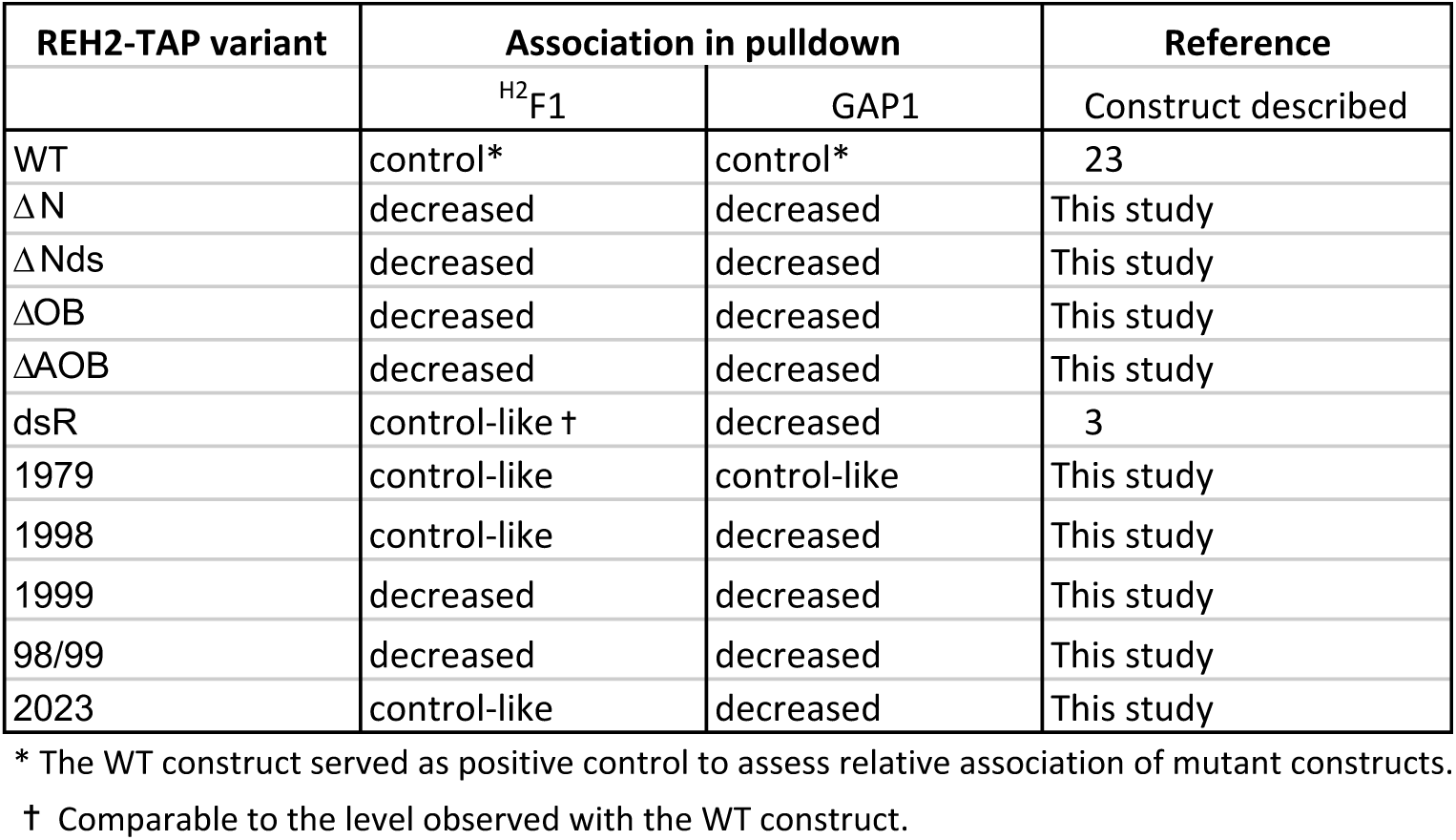
Association of REH2 constructs with ^H2^F1 and GAP1.

| REH2-TAP variant | Association in pulldown |  | Reference |
| --- | --- | --- | --- |
|  | <sup>H2</sup>F1 | GAP1 | Construct described |
| WT | control* | control* | 23 |
| ΔN | decreased | decreased | This study |
| ΔNds | decreased | decreased | This study |
| ΔOB | decreased | decreased | This study |
| ΔAOB | decreased | decreased | This study |
| dsR | control-like † | decreased | 3 |
| 1979 | control-like | control-like | This study |
| 1998 | control-like | decreased | This study |
| 1999 | decreased | decreased | This study |
| 98/99 | decreased | decreased | This study |
| 2023 | control-like | decreased | This study |

\* The WT construct served as positive control to assess relative association of mutant constructs.
† Comparable to the level observed with the WT construct.

### Analysis of ^H2^F1 mutants *in vivo*

To analyze specific features of the ^H2^F1 zinc finger protein in trypanosomes, we expressed a Tet-inducible TAP tagged ^H2^F1 WT construct and examined its association with endogenous REH2, GAP1 and REL1 proteins. REL1 is a subunit of the editing enzyme RECC. We found that the tagged ^H2^F1 WT was able to reconstitute interactions of the native ^H2^F1 protein in trypanosomes (Fig. 3). Notably, expression of the ectopic ^H2^F1 WT increased the steady-state level of endogenous REH2 compared to a control lane without tetracycline (Fig. 3A, lanes 1-2; Fig. 3B). As a control to mark the position of REH2 we used a reported RNAi-based knockdown of ^H2^F1 that decreases the level of endogenous REH2 (Fig. 3A, lanes 5-7; Fig. 3B)(4). The contrasting effects of the upregulation and downregulation of ^H2^F1 on the endogenous REH2 are consistent with the idea that ^H2^F1 stabilizes REH2 *in vivo*. We examined the distribution of REH2 and ^H2^F1, and canonical proteins in RESC and RECC, in sedimentation analyses of mitochondrial extract (S5 Fig). REH2 is heterodispersed as shown before (23) and overlaps with ^H2^F1 in the gradient. However, the relative abundance of these proteins at high and low densities differs substantially. It is possible that the association between REH2 and ^H2^F1 or their stoichiometry in REH2C varies *in vivo*. Additional studies will be necessary to examine these possibilities.

**Fig 3.**
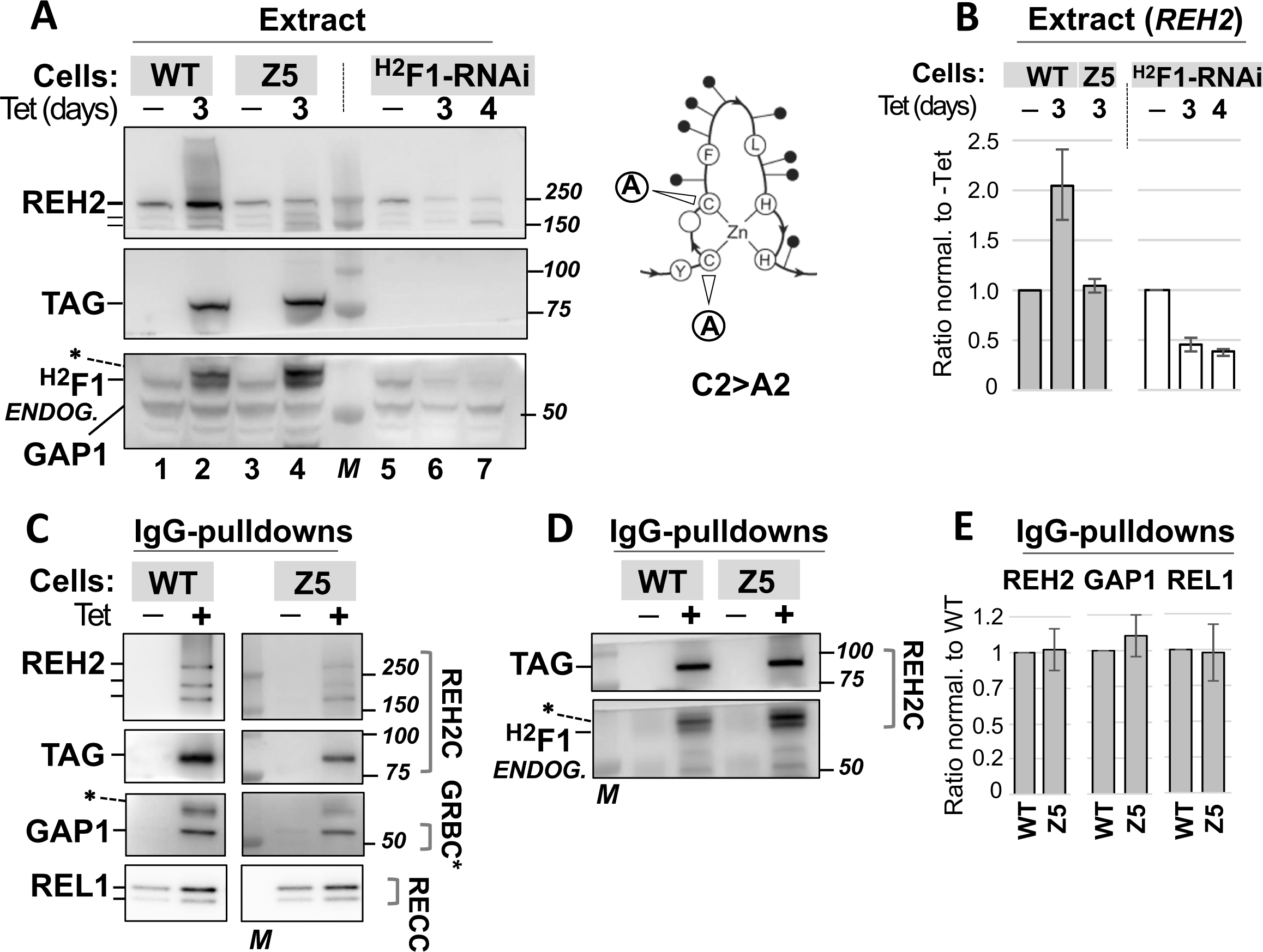
Effect of expressing ^H2^F1 WT or Znf5 C2>A2 on the REH2 steady-state abundance. **(A)** Western blots of extracts with tagged ^H2^F1 WT (lanes 1-2) or Z5 C2>A2 (Z5; lanes 3-4). Control assays include lanes without Tet-induction. Control lanes with a reported ^H2^F1-RNAi construct mark the position of the endogenous REH2 and ^H2^F1. The RNAi lanes show the induced decrease of ^H2^F1 and concurrent fragmentation of REH2 full-length. Hash marks indicate full-length and two prominent fragments of REH2 (4). GAP1 was used as a loading control in all assays in this panel. Molecular size markers (M). A typical C2H2 zinc finger domain fold is depicted with core cysteine substitutions C2>A2 (arrows), other conserved residues, and variable residues (black circles). **(B)** Charts of the relative levels of endogenous REH2 full-length in the extracts +/-1 SD, *n*=3. Induced/uninduced ratios of REH2 for each construct were further normalized to GAP1. **(C-D)** Western blots of IgG pulldowns of WT and Z5 constructs showing association with endogenous REH2, GAP1, REL1, and endogenous ^H2^F1. REL1 is examined by radiolabeled self-adenylation. Other proteins are examined as in panel A. Non-specific species (*) may represent fragments of the tagged protein that react with the antibodies. **(E)** Charts of the relative levels of REH2, GAP1 and REL1 in the pulldowns +/-1 SD, *n*=3. The ratios (protein/tagged-bait) in the Z5 mutant were normalized to ratios in the ^H2^F1 WT construct.

To determine if the stabilizing effect of ^H2^F1 requires one or more zinc finger (Znf) domains, we introduced amino acid substitutions that either disrupt the canonical fold of the domain (by replacing the canonical residues) or eliminate the positive charge (by replacing various basic residues with neutral residues) in individual fingers of the ectopic construct. The examined ^H2^F1 mutants are summarized in Table 2. The location of the fingers and the examined substitutions (including between 2 and 6 basic residues per finger) in the full sequence of ^H2^F1 are shown in S6 Fig. All ^H2^F1 mutants examined were directly compared and normalized to a WT construct used as control in all panels. This enabled the separation of mutants in multiple panels.

**Table 2.** ^H2^F1 mutants with reduced ability to stabilize REH2 in vivo.

| <sup>H2</sup>F1-TAP variant | Endogenous REH2<br>*Steady-state level* | Figures using this<br>construct |
| --- | --- | --- |
| WT | Increased* | Figs. 3, 4 and 5 |
| Znf5 C2>A2 | no change<sup>†</sup> | Fig. 3 |
| Znf1 R/K>A | no change | Fig. 4 |
| Znf2 R/K>A | no change | Fig. 4 |
| Znf3 R/K>A | no change | Fig. 4 |
| Znf4 R/K>A | no change | Fig. 5 |
| Znf5 R/K>A | no change | Fig. 4 |
| ΔN | no change | Fig. 5 |
| ΔC | no change | Fig. 5 |

\* <sup>H2</sup>F1-WT ectopic expression increased the level of endogenous REH2
† No change (i.e., level within standard deviation for at least two biological replicates)

#### Canonical cysteine residues

We changed two cysteines to two alanines (C2>A2) to disrupt the native conformation of the predicted zinc fingers (Fig. 3A). We began by disrupting Znf5 because of its significant homology to a reported dsRNA-bound C2H2 zinc finger domain structure (4, 14, 24). Mutants are referred to in the text by the acronym in the general form Znf*X*. A shorter version of the acronym is used in the figure annotations. The resulting mutant protein Znf5 C2>A2 did not induce an evident increase in the steady-state level of endogenous REH2 as we found after expressing the ^H2^F1 WT construct. This was evident in direct comparisons of extracts that contain similar levels of ectopic ^H2^F1 protein, WT or Znf5 C2>A2 (Fig. 3A, lanes 1-4; Fig. 3B). To examine whether the lack of Znf5 affected the association of ^H2^F1 with other editing components, we performed IgG pulldown assays of tagged ^H2^F1. These assays showed that the ^H2^F1 WT and Znf5 C2>A2 variants similarly associated with endogenous helicase REH2, GAP1 and REL1 ligase (Figs. 3C, 3E). Notably, pulldowns of both tagged ^H2^F1 WT and Znf5 C2>A2, contain endogenous ^H2^F1 (Fig. 3D). The above data suggest that Znf5 is not essential for the normal association of ^H2^F1 with other editing components. We cannot rule out that the endogenous ^H2^F1 is “saving” the examined associations with the mutant ^H2^F1.

We also attempted to generate a ^H2^F1 variant bearing a triple C2>A2 substitution in Znf6, Znf7 and Znf8. However, expression of this construct was unstable and eventually lost in culture. Together, the observations above suggest that ^H2^F1 may require a full complement of zinc fingers in their native conformation to stabilize REH2 but not to associate with the examined editing components.

#### Variable basic residues

Because the native conformation of the zinc fingers seems required for ^H2^F1 to stabilize REH2 we introduced mutations that decrease the positive charge in the zinc fingers yet likely to retain native conformation (25). We expressed ^H2^F1 variants to test the role of several basic residues around the core residues that are mostly located between the second cysteine and the first histidine (Fig. 4A) (26). Specifically, we introduced R/K>A substitutions (between 2 and 6 per finger) to decrease the positive charge in the zinc fingers and to remove potential charge-charge interactions with REH2 or associated RNAs. As above, we use in the text the acronym in the general form Znf*X* but a shorter version of the acronym in the figures. Our functional analyses of Znf1, Znf2, Znf3 or Znf5 R/K>A variants showed that the loss of positive charge in any of these zinc fingers reduced the ability of ^H2^F1 to induce the accumulation of intact endogenous REH2 in extracts (Figs. 4B-C). However, these mutations did not prevent the association of ^H2^F1 with REH2, GAP1 and REL1 (Figs. 4D-F). These expressed ^H2^F1 variants also co-purified with endogenous ^H2^F1 at a level comparable to the WT construct (Figs. 4D, 4F). The data above indicated that the wildtype and R/K>A variant proteins associated similarly with the examined editing components. Overall, ^H2^F1 may require that most zinc fingers retain their native conformation and positive charge to efficiently increase the stability of REH2. However, the loss of the native conformation or positive charge in individual fingers does not preclude the assembly of ^H2^F1 in the REH2C, or the association of this subcomplex with the examined editing components. Detection of endogenous ^H2^F1 in our pulldowns suggest the presence of ^H2^F1 multimers. Pulldowns treated with an RNaseA/T1 mixture suggest that the association between tagged ^H2^F1 and endogenous ^H2^F1 is RNase-resistant (S7 Fig.). We note the possibility that the tested ^H2^F1 mutants may exhibit defects in the examined associations and that these defects are masked by the endogenous ^H2^F1. However, the reduced stabilizing property of the mutants was not saved by the endogenous ^H2^F1. Further studies using isolated recombinant proteins are needed to establish direct protein contacts between ^H2^F1 monomers as was shown between REH2 and ^H2^F1 (4).

**Fig. 4.**
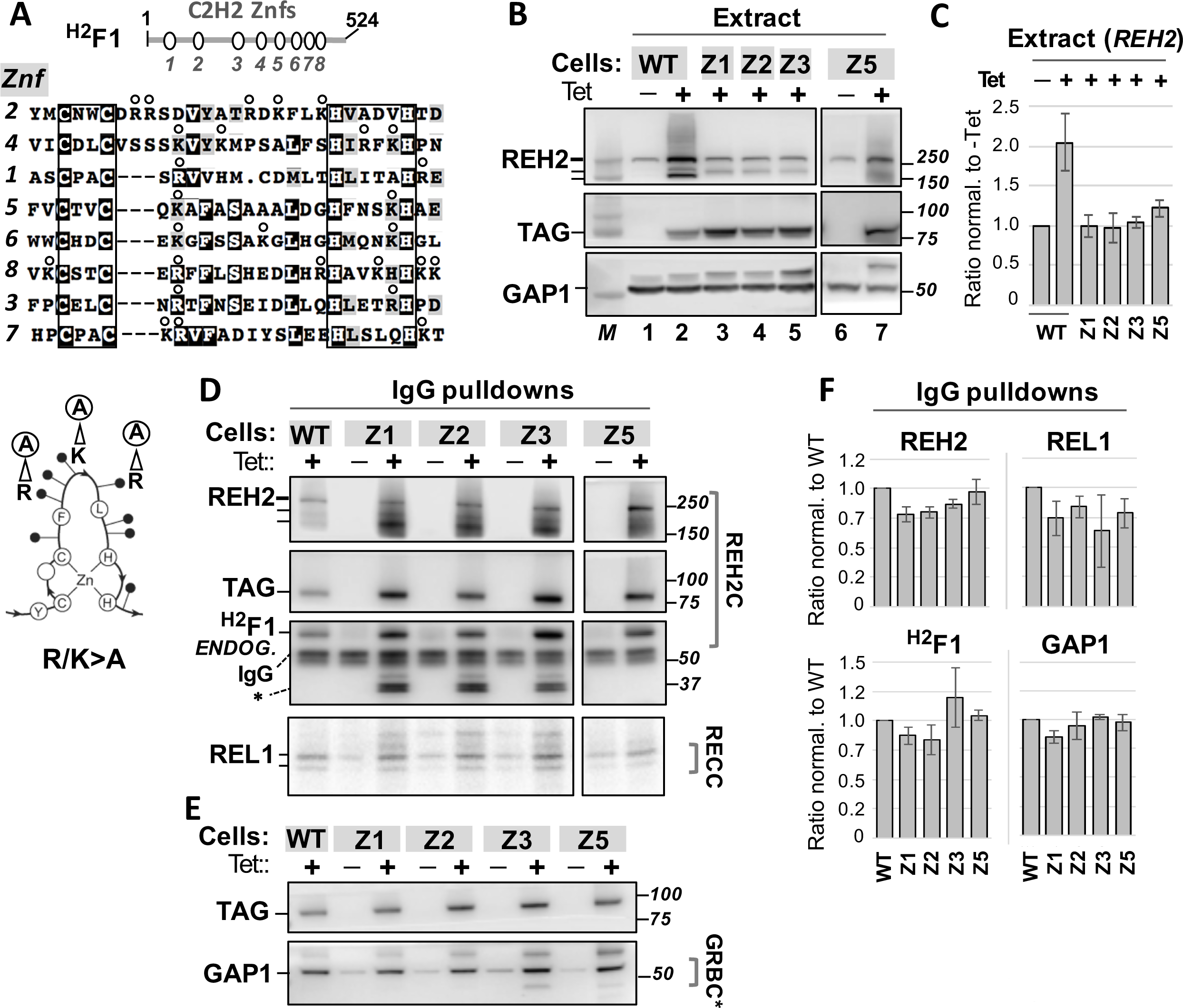
*In vivo* analyses of ^H2^F1 variants with reduced positive charge in Znf1, Znf2, Znf3 or Znf5 (variable residues R/K>. **A)**. Figure annotations use a shorter acronym of each mutant in the form: Z*X,* as in Fig. 3. **(A)** Cartoon of ^H2^F1 (524 residues). Multi-sequence alignment of the eight C2H2 zinc fingers, indicating the core cysteines and histidines (boxes) and R/K>A substitutions of variable residues (between 2 and 6) in each finger (open circles). The C2H2 zinc finger domain fold is depicted as in Fig. 3. In this case, the arrows illustrate three variable residue substitutions R/K>A. **(B)** Western blots of extracts as in Fig. 3A of induced tagged ^H2^F1 WT and Z1, Z2, Z3 and Z5 R/K>A mutants or uninduced control. GAP1 was used as a loading control (lanes 1-7). **(C)** Chart of steady-state level of endogenous REH2 full-length in extracts with each construct as in Fig. 3B. **(D-E)** Western blots and self-adenylation reactions of IgG pulldowns of ^H2^F1 WT and Z1, Z2, Z3 and Z5 R/K>A mutants showing association with endogenous REH2, ^H2^F1, REL1 and GAP1. Non-specific species in pulldowns (*) are indicated. **(F)** Charts of the relative level of associated endogenous proteins in the pulldowns +/-1 SD, *n=2* or *n*=3, as in Fig. 3. The ratios (protein/tagged-bait) in the mutants were normalized to ratios in the ^H2^F1 WT construct.

#### N- and C-truncations

Because mutagenesis of the tested Znf domains did not prevent the association of ^H2^F1 with other editing components *in vivo*, we asked if fragments of ^H2^F1 retain these interactions. To test this question, we expressed N-or C-terminal fragments including Znf1-to-Znf3 (termed ΔC) or Znf4-to-Znf8 (termed ΔN) that span 49% and 51% of the ^H2^F1 polypeptide chain, respectively (Fig. 5A, bottom). These two fragments exhibited minimal or no association with the examined endogenous REH2, ^H2^F1, or REL1 ligase (Figs. 5A). These mutants were directly compared with the ^H2^F1 WT and Znf4 R/K>A constructs. We note that while Znf4 R/K>A was examined separately from the R/K>A variants in Fig. 4, all R/K>A mutants are normalized to the ^H2^F1 WT construct and uniduced lane that were included in each gel. As with other R/K>A mutants, Znf4 R/K>A did not cause an evident increase in the steady-state level of endogenous REH2 in extracts as we found with the ^H2^F1 WT construct. The relative level of endogenous REH2 was 1.0 +/-10% in the induced extract with Znf R/K>A normalized to the uninduced control. The Znf4 R/K>A and other Znf R/K>A mutations in the current study retained all tested interactions that are observed with the ^H2^F1 WT control. However, the ΔN and ΔC constructs failed to establish basic interactions with the tested editing components. Because the native REH2C subcomplex carries mRNA (4), we wondered if the ^H2^F1 ΔN and ΔC constructs also fail to co-purify with mRNA. To this end, we quantitated the relative content of mitochondrial transcripts in the pulldowns of the WT, ΔN, and ΔC constructs in biological replicate experiments that were performed in triplicate (Fig. 5B). The examined mitochondrial transcripts include examples of unedited, edited, and never-edited mRNA (ND4), and ribosomal RNA 9S. We also examined cytosolic transcripts (Tubulin, 1400 and 1390) to score non-specific RNA interactions of editing subcomplexes as in previous pulldown studies (5). Relative to the ^H2^F1 WT control used for normalization, both ΔN and ΔC pulldowns exhibited a dramatic loss in bound mitochondrial mRNAs (Fig. 5B, upper and middle panels). Interestingly, the level of cytosolic mRNAs was similar in the ΔN and WT proteins but dramatically less in the ΔC variant. The latter variant lost most examined RNA interactions: specific and non-specific. An exception may be mitochondrial RNA 9S. We also compared the relative impact of the ΔN and ΔC deletions with a more discrete alteration in Znf4 R/K>A (Fig. 5B, lower panel). The level of mitochondrial RNAs in the Znf4 pulldown was similar to that of the ^H2^F1 WT control. However, the level of cytosolic transcripts decreased ∼70 fold in Znf4 R/K>A. The independent pulldowns and RNA quantitation in three equivalent experiments showed consistent results in the protein interactions and RNA content of these ^H2^F1 constructs. The above data indicated that the tested ^H2^F1 fragments appear to have lost the normal direct interaction of ^H2^F1 WT with REH2 in REH2C, and the normal RNA-dependent association of ^H2^F1 WT with other examined proteins in the editing apparatus. Quantitation of the examined transcripts in the input extracts (including those used in the pulldowns in Figs. 5A-B) confirmed that the observed dramatic loss in RNA association with ΔN and ΔC constructs was not due to large changes in the transcript level at steady state (S8 Fig.). The ^H2^F1 ΔN construct used a heterologous mitochondrial leader sequence (see methods and S1 Table). A comparison of ^H2^F1 ΔN and ΔC in whole-cell and enriched-mitochondrial extracts suggest that ^H2^F1 ΔN localizes to mitochondria efficiently. However, the import or retention of ^H2^F1 ΔC into mitochondria is relatively compromised. At least some import of the ^H2^F1 ΔC construct may occur in comparison with a control cytosolic marker in whole-cell and enriched-mitochondrial extracts (S2 Fig.). Apparent fragmentation of ^H2^F1 ΔC (TAP tag panel in S2 Fig.) suggests that this construct is also partially unstable. These results indicate that features in the N terminus of ^H2^F1 are required for efficient ^H2^F1 association with mitochondrial editing proteins and transcripts in the pulldowns. The C-terminal truncation affected the ^H2^F1 association with all examined transcripts (including cytosolic) and somehow also its mitochondrial import and stability. The expressed ^H2^F1 ΔC construct may not entirely reflect the ability of the N-terminal half of ^H2^F1 to bind editing proteins and transcripts due to the reduced localization and stability of this construct. Finally, the more discrete alteration in Znf4 R/K>A may have caused a relatively mild decrease in RNA affinity that mostly affected weak, non-specific interactions, e.g., by cytosolic RNA in the pulldowns of ^H2^F1.

**Figure 5.**
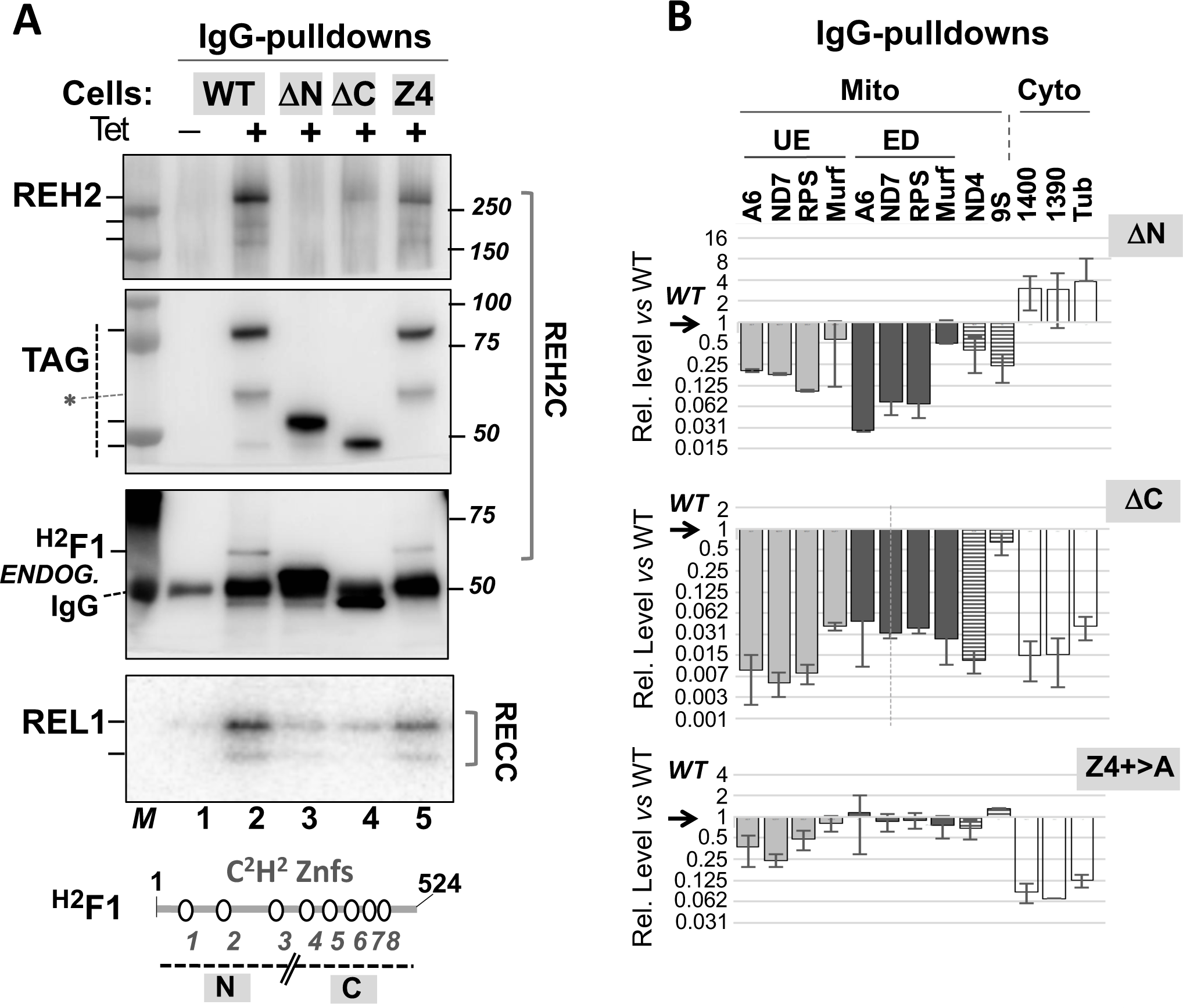
*In vivo* analyses of ^H2^F1 terminal deletion and Znf4 R/K>A mutants. **(A)** Western blots of IgG pulldowns of N and C truncation mutants (ΔN and ΔC) that removed fingers 1-to-3 and fingers 4-to-8, respectively, and the Z4 R/K>A variant. Association of the constructs with endogenous REH2, ^H2^F1 and REL1 was examined. Biological replicates of these assays suggest a similar quality and loading of samples in the pulldowns. Cartoon of ^H2^F1 with zinc fingers and a line marking the N and C truncations. Non-specific species in pulldowns (*) are indicated. **(B)** qRT-PCR quantitation of transcripts in the pulldowns in panel A. dCq of IgG pulldowns of tagged protein (induced extract) normalized to a mock pulldown (uninduced extract) [dCq = 2^(test Cq – mock Cq)^] for each construct. The mutant proteins were then normalized to the tagged WT protein. All values took into account the recovery of tagged protein in each pulldown in panel A. Three independent biological replicates with Cq average values and one standard deviation (+/-1 SD, *n*=3) were plotted. Non-specifically associated cytosolic transcripts differed substantially in pulldowns of different mutants, so these transcripts were not used as reference. Mitochondrial unedited (UE) and edited (ED) and cytosolic RNA transcripts are indicated. RPS, Murf, and 9S are abbreviations for RPS12, Murf2, and 9S rRNA, respectively. 1390 and 1400 are abbreviations for the cytosolic HGPRT isoforms Tb927.10.1390 and Tb927.10.1400, respectively. Tub is tubulin. Other short names are standard.

### Analysis of recombinant REH2 *in vitro*

The conserved predicted catalytic domains in REH2, the unwinding activity in pulldowns of REH2 or ^H2^F1 from extracts, and the loss of this activity in the pulldowns after mutating catalytic amino acids in REH2, indicate that REH2 in the context of its native subcomplex functions as an RNA helicase (3, 4, 23). However, the unwinding activity of the isolated REH2 has not been formally established. We have now generated a his-tagged recombinant full-length rREH2 polypeptide (240 kDa) without its mitochondrial leader sequence (see the methods section) that supports robust ATP-dependent unwinding of a synthetic dsRNA substrate *in vitro.* This polypeptide is largely intact in most preparations (Fig. 6A). The dsRNA substrate in our assay is a mimic of the native A6 editing system formed by pre-annealing of a 3’ fragment of the pre-mRNA A6 with a cognate gRNA. The synthetic A6 mRNA/gRNA pair supports a full round of U-deletion editing in an *in vitro* assay with cell extracts and purified RECC enzyme (27, 28). The unwinding activity of the recombinant enzyme increased in a titration with increasing concentrations of rREH2, and also in a time course up to 30 min when most of the annealed radiolabeled RNA was unwound (Figs. 6B-C). An upshift of radiolabeled RNA in a ribonucleoprotein complex (RNP) at the highest tested concentrations of rREH2 (12 pmoles) in the titration (Fig. 6B) may be due to the known RNA binding activity of the native REH2 established by UV crosslinking (5, 23). The radiolabeled strand (gRNA) alone or in heat-treated controls of the dsRNA substrate exhibit the same gel mobility as the helicase-unwound gRNA (Figs. 6C-D). In summary, the isolated 240 kDa recombinant REH2 is a *bona fide* RNA helicase enzyme *in vitro*, and further supports the idea that REH2 may remodel RNA or its RNPs in editing.

**Figure 6.**
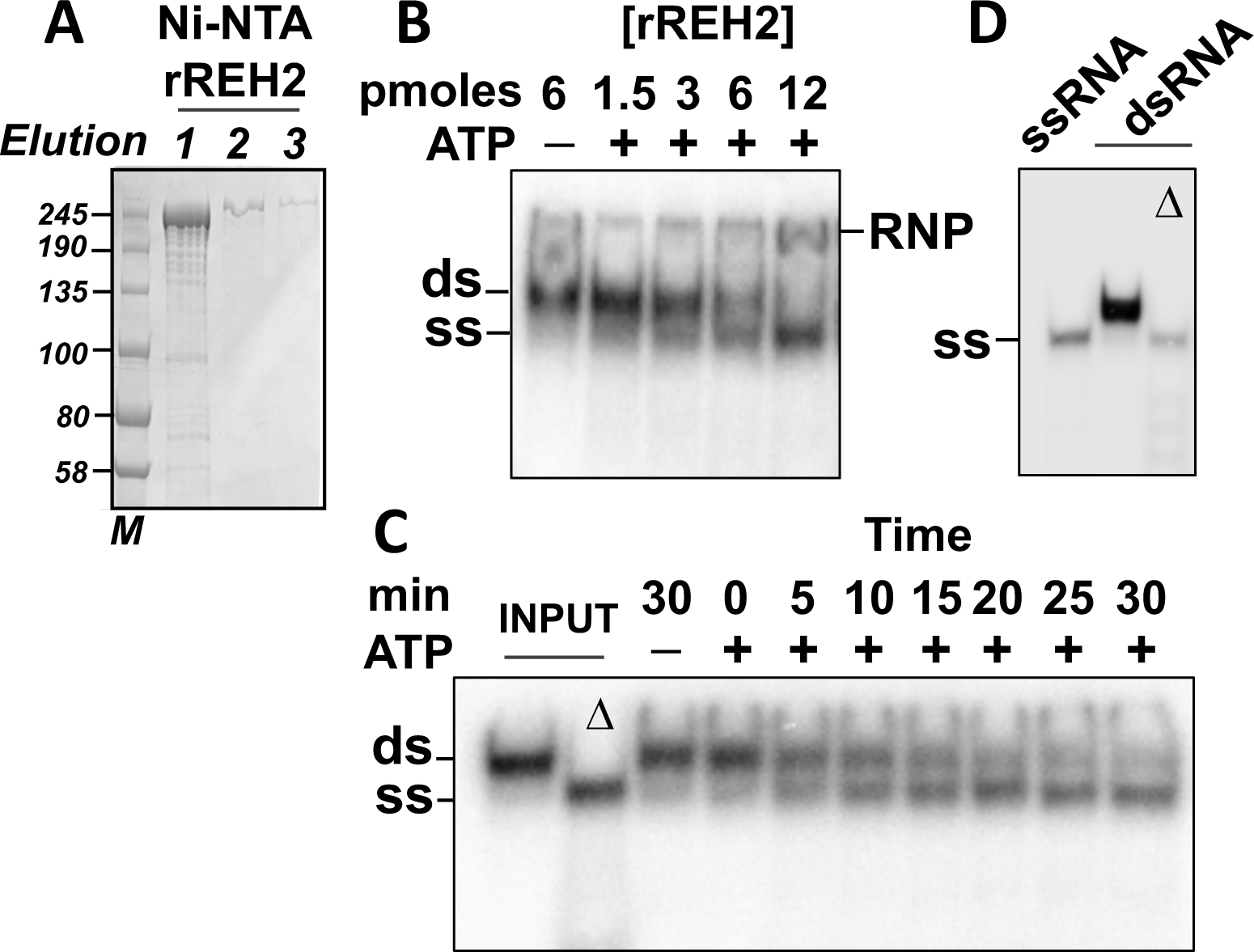
Recombinant rREH2 is a catalytically active RNA helicase. **(A)** Purified rREH2[30-2167] samples in a denaturing gel stained with Coomassie. NTA-Ni elution fractions 1-to-3 are shown with molecular size markers (kDa). **(B)** Unwinding assays in 10 μL reaction mixtures with increasing concentrations of rREH2[30-2167] +/- ATP. An RNP complex (RNP) forms at the highest enzyme concentration tested; **(C)** Unwinding assays at increasing incubation times with rREH2[30-2167] +/- ATP using the standard reaction mixture described in the material and methods section. The standard reaction used a molar excess of enzyme over substrate. The dsRNA (ds) was assembled by pre-annealing of synthetic transcript that mimics a 3’ fragment of the *T. brucei* A6 pre-edited mRNA and a radiolabeled cognate guide RNA (27). Radiolabeled unwound ssRNA (ss) is indicated. Input dsRNA was heat denatured (Δ) as control. The assays in this panel were treated with proteinase K to remove any RNP that may accumulate (see methods). **(D)** Additional controls showing that the starting radiolabeled ssRNA used to generate the dsRNA substrate in the assay and the unwound radiolabeled ssRNA in the heat-denatured (Δ) dsRNA (in panel C) have the same gel mobility.

## Discussion

In the current study, we examined over 20 constructs of REH2 and ^H2^F1 (Tables 1 and 2) to identify features in these editing proteins that impact the stability of REH2, the assembly of the REH2C RNP, and its RNA-mediated coupling with other components of the RNA editing holoenzyme (or RNA holo-editosome). We also formally established that the isolated REH2 is a *bona fide* RNA helicase enzyme.

The mRNA editing *trans* factors core enzyme RECC and accessory RESC and REH2C exhibit transient and stable contacts with each other via RNA. Variants of RECC and the RESC modules, GRBC and REMC, are also emerging in studies by different labs (reviewed in 1). These variants imply a dynamic nature of holo-editosomes that may be important in editing control and needs further characterization. The helicase complex REH2C exhibits transient and stable contacts with variants of RESC that minimally differ in the relative content of MRB3010 (in the GRBC module). This study focused on stable interactions by REH2C that can be readily examined. Prior studies *in vivo* showed that inactivating mutations in the helicase ATP binding site or the dsRBD2 domain dissociate REH2 from mRNA and RESC components, suggesting a general helicase detachment from the editing apparatus (3). ^H2^F1 RNAi dissociates the helicase from canonical proteins in the RESC modules. RNAi knockdown of either ^H2^F1 or REH2 also inhibited the editing associated with a RESC variant that is particularly active and transiently interacts with REH2C (3, 4). These and other observations indicate that transient and stable contacts in dynamic holo-editosomes including those involving REH2C are important.

A systematic mutagenesis in the current study identified features affecting the assembly of the REH2C RNP, or its association with other editing components *in vivo*.

### REH2 mutants *in vivo*

The NTR truncations examined here (>1000 residues each), ΔN and ΔNds, caused a major loss in ^H2^F1 binding and RNA-mediated association with GAP1. The only difference between the ΔN and ΔNds constructs is that the latter lacks the ∼70 amino acid dsRBD2 domain. As mentioned above, the dsRBD2 promotes the addition of REH2 into holo-editosomes (3). Also, the dsRBD2 in the related helicase MLE in flies may provide a major RNA-binding surface (29, 30). The tested truncations suggest that other N-terminal features besides dsRBD2 may contribute to normal interactions by REH2 *in vivo*. These truncations may also introduce topological changes that alter the normal REH2 interactions but do not prevent localization of the constructs in enriched-mitochondrial extracts (S2 Fig.). Conformational changes were proposed to modulate interactions by other eukayotic DExH/RHA helicases (18, 31). Overall, the tested NTR truncations may indicate the presence of unidentified protein features or global changes in conformation that affect the REH2 interactions. Further studies will be needed to resolve these possibilities. However, relatively short CTR truncations and point mutations that affected the REH2 interactions with other editing components are less likely to cause major conformation changes in REH2.

We have speculated that the conserved OB fold in DExH/RHA subfamily helicases may share functional residues in kinetoplastid RNA editing and other RNA processes (14). Our examined C-terminal truncations included most of the CTR (260 residues in ΔOB) or elements beyond the OB fold that may be specific of REH2 (123 residues in ΔAOB). We examined R2023 in REH2 that corresponds to R704 in helicase Prp43p in yeast spliceosomes (16) (S3 Fig.). R2023 should be located in a critical loop, β4-β5, in the REH2 OB fold. In yeast, R704 is poised to interact with the RNA substrate entering the helicase cavity, and mutation of this residue inhibits the RNA binding and RNA-mediated activation of Prp43p (16). We also examined R1979 in REH2, which is not conserved in Prp43p. In trypanosomes, the R2023A substitution had a moderate effect on the stable RNA-mediated association of REH2C with GAP1. However, R1979A did not evidently affect this interaction. Neither R1979A or R2023A seem to significantly affect the REH2 direct interaction with ^H2^F1.

Similarly, H1998 and R1999 in REH2 were compared to the equivalent H1032 and K1033 positions in the RNA helicase MLE in flies. The later residues in MLE bind specific uridylates in U-rich roX boxes of the cognate roX RNA. H1032 and K1033 are also needed for the chromosomal localization of MLE in the *Drosophila* dosage compensation complex (18). The H1998E substitution had a moderate effect on the REH2 association with GAP1 but had little or no effect on the interaction with ^H2^F1. In contrast, the adjacent substitution R1999E affected both the direct REH2 interaction with ^H2^F1 and the RNA-based REH2 association with GAP1. These results are consistent with the idea that ^H2^F1 is an adaptor protein of REH2 (4), and that structure-function correlations may be traceable to specific conserved amino acids in REH2 and related helicases in highly-diverged species (14). This is also in line with the idea that the OB fold in DExH/RHA helicases, including REH2, is a conserved regulatory domain.

### ^H2^F1 mutants *in vivo*

^H2^F1 has eight predicted C2H2 Znf domains that cover over ∼45% of the polypeptide sequence. Apart from the Znf domains, ^H2^F1 has no evident sequence or structural conservation outside of the kinetoplastids. ^H2^F1 is an adaptor protein that brings the REH2 to the editing machinery (4) and it also stabilizes REH2. That is, the induced downregulation of ^H2^F1 caused fragmentation of REH2 (4), and we showed here that the converse upregulation of ^H2^F1 increased the REH2 level *in vivo*. ^H2^F1 may require most of its zinc fingers to stabilize REH2 because loss of the core structure or positive charge in individual fingers compromised this function. It is feasible that the stabilizing effect of ^H2^F1 involves substantial changes in the helicase conformation and coordination of its zinc fingers. Notably, all examined ^H2^F1 substitution-mutants were able to associate with REH2, GAP1, and REL1. This suggested that the basic fold or positive charge of the fingers is not essential for the ^H2^F1 association with other editing components. However, N-or C-terminal truncation of nearly half the size of ^H2^F1 hampered its normal interactions with the examined editing components. The ^H2^F1 ΔN accumulated in enriched-mitochondrial extracts but the ^H2^F1 ΔC localization in mitochondria seemed relatively compromised. Somehow the C-terminal truncation was particularly deleterious to ^H2^F1. Additional studies are needed to pinpoint specific determinants for ^H2^F1 direct binding to REH2 or its RNA-mediated contacts with other components.

Besides the REH2•^H2^F1 system, other DExH/RHA helicase•Znf protein pairs have been characterized (14). In the DXH30•ZAP system in antivirus protection in humans, the antiviral protein ZAP has four Znf motifs that participate in viral RNA recognition (32, 33). All four fingers lie on a positively charged surface in a crystal structure of ZAP (25) and mutation of basic residues in the fingers inhibited the ZAP antiviral function (33). The ZAP studies imply that stabilization of REH2 by ^H2^F1 could involve a precise topology of its zinc fingers, and that the ^H2^F1 fingers may participate in RNA binding. Notably, recombinant ZAP is multimeric (25). If ^H2^F1 dimerizes, a single REH2C RNP would include 16 zinc fingers. The co-purification of tagged and endogenous ^H2^F1 from RNase-treated extracts imply direct protein contacts between the ^H2^F1 copies (S7 Fig.). However, analyses of recombinant ^H2^F1 WT and mutants are necessary to establish the potential of ^H2^F1 to multimerize and bind RNA, and to define the role of its fingers. The RNA content in pulldowns of the ^H2^F1 Znf4 variant with four R/K>A substitutions suggested minimal changes in association with target RNAs in mitochondria but a significance loss in non-specific contacts with unrelated transcripts. This suggests that subtle changes in RNA affinity are more likely to affect weak non-specific RNA contacts. The general loss of transcripts examined in the pulldowns of the tested ^H2^F1 fragments also suggests that efficient RNA association by the REH2C complex *in vivo* involves at least the majority of zinc fingers in ^H2^F1. We note that the transcripts examined may associate with either ^H2^F1, REH2 or both proteins in the native REH2C RNP.

### REH2 may function as an RNA helicase in kinetoplastid RNA editing

Prior studies suggested that REH2 is a functional helicase in RNA editing. REH2 pulldowns from GAP1-depleted extracts exhibit RNA unwinding activity (4), and the mutation of catalytic residues in the predicted RecA1 domain inhibits the unwinding activity in pulldowns of REH2 (23). We have now established that the isolated full-length recombinant REH2 (240 kda) is a *bona fide* RNA helicase enzyme. This further indicates that REH2, the only DExH/RHA helicase in kinetoplastid holo-editosomes provides an ATP-dependent unwinding activity in RNA editing. Additional studies are needed to determine if ^H2^F1 affects the activity of REH2. Besides REH2, the editing apparatus also requires the DEAD-box RNA helicase REH1 (12, 13). REH1 is not part of the RNPs studied here and seems to lack a direct binding partner. A transient RNA-mediated interaction of REH1 with other editing proteins has been detected (23, 34). REH1 and REH2 are functionally different. REH2 is needed for editing within single blocks, including the first “initiating” block (3). In contrast, REH1 is needed for editing of two or more blocks (i.e., by overlapping gRNAs) but has no effect on the first block, suggesting a role in gRNA exchange (13). Both RNA helicases may promote critical remodeling of RNA or RNP structure. Our ongoing RNA-seq analyses may provide additional functional insights concerning REH2C (Kumar et al., unpublished).

### Possible roles of editing complex variants

The editing apparatus much like the splicing and transcription molecular machines may exhibit dynamic phases of assembly and function. The RECC enzyme has multiple isoforms that include several RNase III (RIII)-like protein subunits. These RIII proteins have the potential to form a large number of binary combinations that could modulate substrate recognition (35–37). Also, natural GRBC and REMC isoforms and genetically-induced changes in their protein/RNA organization are emerging. A recurring theme is the variable interaction between the essential GAP1/2 heterotetramer and MRB3010 (GRBC6) (1, 6, 38) that is potentially regulatory. For example, GRBC variants in pulldowns of MRB3010 or REH2 differ in relative content of MRB3010 and editing level in associated mRNAs (3, 5). Also, RNAi of REH2 or ^H2^F1 caused a reduced editing in mRNAs in the MRB3010 pulldowns (3, 4). REH2 and ^H2^F1 features, including those examined here, may affect specific steps in the editing mechanism and we are currently addressing this.

## Materials and Methods

### Cell Culture

*T. brucei* Lister strain 427 29-13 procyclic was grown axenically in log phase in SDM79 medium (39) and harvested at a cell density of 1–3×10^7^ cells/mL. All transgenic cell lines were induced with tetracycline at 1μg/mL. The REH2 WT-TAP and ^H2^F1 RNAi cell lines that were used as controls were generated in our lab (3, 4). We have seen that the TAP tag in REH2, including the WT construct, seems to slow the growth of trypanosomes after several days of culture (23). This and presumably incomplete cloning after transfection may account for the uneven expression levels in extracts from independent subcultures. To partially control for this issue, most plasmid constructs used to generate the current data figures were made to use puromycin selection. Most transfections used tetracycline-screened HyClone Fetal Bovine Serum (FBS) (SH30070.03T; GE Healthcare). This enabled a faster cell recovery after transfection than with constructs using phleomycin selection that were originally used in cell lines and appeared somewhat less stable during subculture. Regardless of this issue, independent analyses of protein associations were consistent with either puromycin or phleomycin constructs.

### DNA Constructs

All overexpression and RNAi studies used inducible plasmid constructs. The constructs for ^H2^F1 RNAi and the REH2 dsRBD2 mutant K1078A/A1086D were previously reported (3, 4). N-terminal deletion constructs for REH2 (ΔN and ΔNds) and ^H2^F1 (ΔN) were prepared using the In-Fusion® HD Cloning Kit (639648; Clontech). Two PCR amplicons were prepared in each case using CloneAmp™ HiFi PCR Premix (638500; Clontech) and were then gel-eluted using NucleoSpin® Gel and PCR Clean-up (740609.10; Clontech). One amplicon encodes the first 34 amino acids of REH2 (primers: F-1503, R-1522), including the predicted REH2 mitochondrial leader sequence (REH2-MLS) (23), that was used for all N-deletion constructs. The other amplicon included the remaining relevant gene fragment sequence: 3070-6501 bases (REH2 ΔN), 3280-6501 bases (REH2 ΔNds), or 766-1572 bases (^H2^F1 ΔN). Three-piece in-fusion reactions were performed to join the two amplicons and pLEW79-ada-TAP plasmid linearized at *XhoI* and *BamHI* (40). The PCR primers (S1 Table) included 5’ and 3’ terminal homology (15 bases) to enable the recombination-based fusion between the amplicons and the plasmid DNA. The REH2 (ΔAOB and ΔOB) and ^H2^F1 ΔC deletion constructs are also recombination-based fusions of the amplified relevant gene fragment with linearized plasmid DNA. The REH2 constructs R2023A, H1998E, R1999E and H1998E/R1999E, and the ^H2^F1 construct Znf5 2C>2A were created by site-directed mutagenesis in inverse-PCR reactions of the pLEW79-ada-TAP WT plasmid as the template. pLEW79-ada-TAP was modified from the original pLEW79-TAP (40, 41). The resulting linearized mutant constructs were circularized using the In-Fusion HD Cloning Kit. Other ^H2^F1 constructs used synthetic gBlock gene fragments (IDT) with mutations. N-terminal fragments of ^H2^F1 (Znf1, Znf2 and Znf3 R/K>A single finger mutants) replaced the corresponding sequence in the ^H2^F1 WT construct after digestion with *XhoI* and the internal *ApaI* site in ^H2^F1. A C-terminal fragment of ^H2^F1 to generate Znf4R/K>A or Znf5 R/K>A replaced a corresponding sequence in the ^H2^F1 WT construct after digestion with *ApaI* and *BamH1*. The ^H2^F1 WT construct was made by ligation of an amplicon with the full open reading frame into pLEW79-ada-TAP plasmid linearized at *XhoI* and *BamHI.* All constructs were confirmed by DNA sequencing, linearized with *NotI*, and transfected in trypanosomes (39).

### Western Blots

Western blots of REH2, ^H2^F1 and ^H2^F2 (subunits of REH2C), GAP1 (alias GRBC2; subunit of GRBC) and A2 (alias MP42; subunit of RECC) were performed as reported (4, 23). Western blots of the GAP1/GAP2 (alias GRBC2/GRBC1) paralogs simultaneously, and of MRB8170 and MRB6070 (subunits of REMC and MRB6070/1590 assemblies, respectively), were performed as previously described (42, 43). GAP1 alone was examined in the main data figures. Both GAP1/GAP2 were examined in S1 and S5 figures. Peroxidase Anti-peroxidase antibody (P1291; Sigma) diluted to 1:200 (v/v) was used to detect the TAP-tag. EF-1 α1 antibody (sc-21758; Santa Cruz Biot) diluted to 1:200 (v/v) was used as a cytosolic marker. Protein quantitative analyses using Amersham ECL™ western blot detection reagents were performed in an Amersham Imager 600.

### Radioactivity Assays

RNA ligases in the RECC enzyme were radiolabeled by self-adenylation directly on the beads in IgG pulldowns (44). Unwinding assays of REH2 used reported conditions with a few modifications (4) and full-length recombinant His-REH2. The dsRNA substrate in these assays used an A6 mRNA/gRNA pair (23). The A6 pair mimics the endogenous ATPase subunit 6 substrate. The gRNA was transcribed with RNA T7 polymerase from an amplicon (S1 Table) after *DraI* restriction digestion and gel purification of the product. The 61-nt gRNA was 5’-labelled with γ^32^P-ATP using T4 Polynucleotide kinase. The 62-nt mRNA was synthetic (IDT). A mix with 3 pmoles of labelled gRNA and 12 pmoles of mRNA was heated at 95° C for 3 min, and then immediately incubated for 30 min at room temperature in the annealing buffer (10 mM MOPS pH 6.5, 1 mM EDTA, and 50 mM KCl). The annealed RNA hybrids were isolated from native 12% PAGE run at 4°C at 50 V for 120 min in 0.5x Tris-borate-EDTA buffer. Standard 10 μL reaction mixtures in unwinding buffer (40 mM Tris-Cl pH 8.0, 0.5 mM MgCl^2^, 0.01% Nonidet P-40, and 2 mM DTT) were incubated for 30 min at 19°C with 20 cps of hybrid (∼20 fmoles) and 3 µg of recombinant REH2 (∼12 pmoles). Variations to the standard assay including in the concentration of REH2 or the reaction time are indicated in the text. All assays contained 100 fmoles of unlabeled gRNA as competitor to prevent re-annealing of unwound γ^32^P-gRNA. The assays were stopped with an equal volume of 2x helicase reaction stop buffer (50 mM EDTA, 1% SDS, 0.1% bromophenol blue, 0.1% xylene cyanol, and 20% glycerol) and kept on ice for 5 min. The assays in Fig. 6C were also treated with 0.4U of proteinase K (P8107S; NEB), 25 mM EDTA and 0.5% SDS at 22° C for 30 min. The entire reaction was loaded onto a native 12% PAGE and resolved at 25 V for 120 min at 4°C.

### Cell Extracts and Purification of Protein and RNA from IgG Pulldowns

Whole cell extracts as well as mitochondria-enriched (mini-mito) extracts were used in this study. Whole-cell extracts were generated by incubation of cell pellets in MRB-Triton buffer (25 mM Tris-Cl pH 8.0, 10 mM MgOAC, 1 mM EDTA, 10 mM KCl, 5% glycerol, 0.5% Triton X-100) at 4 °C for 45 minutes followed by centrifugation at 15,000g for 30 minutes. Mitochondria-enriched extracts were prepared using a simplified protocol similar to others reported earlier (2, 45). Namely, cell pellets were resuspended into hypotonic DTE buffer (1 mM Tris-HCl pH 8.0, 1 mM EDTA pH 8.0) supplemented with protease inhibitors and disrupted in a Dounce homogenizer. The suspension was passed through a 26-gauge needle as originally described (46). The enriched mitochondrial vesicles were spun down and the cytosolic content in the supernatant discarded. This step improved the recovery of intact REH2 compared to whole-cell extracts. REH2 is more prone to fragmentation than other proteins examined in this study. The following steps were also as in (46). The pellet was resuspended into STM buffer (0.25M Sucrose, 20 mM Tris-HCl pH 8.0, 2 mM MgCl_2_) supplemented with DNase I, RNase-free (EN0523; Fisher Scientific). The reaction was stopped with one volume of STE buffer (0.25 M Sucrose, 20 mM Tris-HCl pH 8.0, 2 mM EDTA pH 8.0). The enriched mitochondria were spun down and washed in STE buffer. The pellet was resuspended into MRB-Triton buffer and incubated in ice for 15 min. The suspension was spun down at 15,000g at 4 °C for 30 min. The supernatant was collected and stored at −80°C. In IgG pulldowns of the extracts, ectopically expressed TAP-REH2 and TAP-^H2^F1 were recovered as reported (4, 23) with some modifications. Approximately 2 mg of protein was mixed with 1x SUPERase·In™ RNase inhibitor (Invitrogen™) and incubated with Dynabeads IgG (11203D; Invitrogen™). The beads were washed five times with 1ml of wash buffer (150 mM NaCl, 1 mM EDTA, 10 mM MgAOc, 0.1% NP-40, and 25 mM Tris pH 8.0). Protein was eluted from the beads with 15 µL of 1x SDS loading buffer at 95 °C for 2 min. RNA was extracted by treating the beads with 4U proteinase K (NEB) for 2 hrs at 55 °C, followed by phenol extraction and ethanol precipitation. For some pulldowns, RNase A/T1 mix (EN0551; Fisher Scientific) was applied at 20 U/mg of protein both in the input extract and again while bound to the beads.

### Quantitative RT-PCR

Isolated RNA from the pulldowns or directly from input extracts was treated with DNase I, RNase-free (EN0523; Thermo Scientific™) prior to cDNA synthesis with the iScript™ Reverse Transcription Supermix (Bio-Rad) as described elsewhere (5). The amplifications mixtures (10 µl) used SsoAdvanced™ Universal SYBR^®^ Green Supermix (Bio-Rad) and reported oligonucleotides for unedited mRNAs, fully edited mRNA, and reference transcripts (47). Diluted samples of the examined cDNA produced a single amplicon during linear amplification under our conditions. The end-point amplicons described here have been gel-isolated, cloned and confirmed by sequencing as previously described (5). Fold-enrichment of mitochondrial and cytosolic transcripts in IgG pulldowns of tagged proteins (induced extracts), relative to a mock pulldown (uninduced extract), was calculated as follows: Fold = 2[^-ddCq^], where ddCq = Cq test IP – Cq mock IP, as in (5). The relative value for each mutant construct was adjusted to the amount of recovered tagged protein, and subsequently normalized to the tagged WT construct (= 1). The plot in Fig. 5B was constructed using data from 3 independent biological replicate experiments, including a total of six amplicons per data point (i.e., two technical replicate amplicons per assay). Two of the independent experiments (including cultures +/-Tet for each mutant) were performed concurrently. The third independent experiment (one set of cultures per mutant +/-Tet) was conducted on a subsequent date. The replicate experiments showed a consistent sample loading in the assays used in our plots. The examined mutant constructs differed in the amount of non-specific association with Tubulin and other cytosolic transcripts that are normally used as internal reference in antibody pulldowns using a common protein bait. The raw Cq values of examined editing transcripts in the pulldowns and the plotted values after normalization indicated a substantial loss of these transcripts in the truncated mutants. qRT-PCR assays of input extract in S8 Fig. were normalized using the following equation: Ratio (reference/target) = 2^[Cq(ref) − Cq(target)]^. This scores the relative difference between the reference and target Cq values. The current plots of input extract used 18S rRNA as reference (S8 Fig.). We obtained similar results with any of these cytosolic transcripts used as a reference: tubulin, 1400 or 1390 (Tb927.10.1400 or Tb927.10.1390, respectively).

### Homology Modeling and Bioinformatics Analysis

The multiple-sequence alignment of REH2 and MLE, and that of the ^H2^F1 zinc finger domains, was done with Clustal Omega (48). Formatting of the fingers alignment was done with BOXSHADE version 3.21 (http:/www.ch.embnet.org/software/BOX_form.html). Sequence alignment of Prp43p, REH2, and other helicases was done using the CDD tool in NCBI (49). The structure-based searches of *T. brucei* REH2 used the program Phyre2 (20). The homology model of REH2 derived from the structure of yeast Prp43p. The inset is a closer view of R2023 and the surround region in the OB-fold. The molecular surface is colored by residue conservation. The image was made using PyMOL version 2.2.

### Recombinant Protein

6xHis-REH2 (amino acid residues 30-2167) was amplified by PCR (S1 Table). The gel-purified product was cloned into the NcoI and BamHI sites in the expression vector pET15b by using the NEBuilder® HiFi DNA Assembly Master Mix (E262, NEB) and the appropriate primers. The ligation reaction mixtures were transformed into chemically competent Omnimax cells from ThermoFisher Scientific. Colonies with the correct insert were identified by single-colony PCR and then amplified for plasmid purification. The sequence of the flanking vector region and the insert was checked by DNA sequencing. The plasmid DNA was transformed into Rosetta2 DE3 cells (Novagen Inc.) and overexpressed in 4 L of Terrific Broth media by supplementing 2% (v/v) ethanol and 2 mM MgCl_2_ at 37 °C. Protein induction with 0.5 mM IPTG was done at a OD_600_ of 0.8, and the expression was continued for 22 hours at 16 °C. The cell pellet was stored at −80°C overnight and was resuspended in lysis buffer (5 mL/g of cell pellet) of 50 mM Tris-HCl pH 8.0, 250 mM NaCl, 1 mM EDTA, and 1 mM DTT along with 10 µg of RNase-free DNase and 15 mg of hen egg white lysozyme. The cells were lysed in an Emulsiflex hydraulic press. The cell debris was pelleted at 39,200 x g for 30 minutes at 4°C. The supernatant with 6xHis-REH2 was batch attached to equilibrated Qiagen Ni-NTA Agarose at 4°C for 16 hrs. The helicase was eluted with 50 mM Tris-HCl pH 8.0, 300 mM NaCl, 1 mM DDT, and 250 mM imidazole. The first elution of the purification was dialyzed into unwinding assay buffer. This material was tested for the presence of the recombinant protein in 6% SDS-PAGE stained with Commassie blue, western blots and antibody pulldowns, and for activity in dsRNA unwinding activity assays *in vitro*.

## Acknowledgements

We thank the Read lab for their gift of antibodies against GAP1, TbRGG2 (alias RGG2), MRB6070, MRB8170, and ^H2^F2. The Afasizhev lab kindly provided antibodies that react against the GAP1/GAP2 paralogs (alias GRBC2/GRBC1) simultaneously. We also thank the Stuart lab for providing the antibodies against A2 (alias MP42). Amna Mushtaq provided detailed comments on the manuscript. Jana Gomez, Elizabeth Alvarez and Melissa Yang helped with the preparation of figures.

## Supporting information

**S1 Figure. Western blot analyses of RESC associated proteins in antibody pulldowns of REH2 and MRB3010**. Immunoprecipitations (IP) from extract with specific antibodies against REH2, MRB3010 or CoxI (mock). **(A)** Western blot analyses examined for the presence of REH2, MRB8170 and MRB6070 proteins. These are markers of REH2C, REMC and a MRB6070/MRB1590-containing subcomplex that associates with RESC, respectively. Extract (Ext) used in the IPs and size markers in kDa (M) are indicated. **(B)** Western blot analyses as in A examining REH2, the GAP1/GAP2 paralogs, and MRB6070. We note that proteins in the IPs are often upshifted relative to the control extract lane. The cause of this upshift is unclear to us but it may be a slight effect on the proteins migration due to the presence of IgG in those samples. IgG in the pulldown lanes is indicated.

**S2 Figure. Ectopically-expressed TAP constructs in whole-cell and enriched-mitochondrial extracts**. Western blots of large-truncation constructs in whole-cell (WC) and enriched-mitochondrial (aka Mini-Mito “MM” extract): (A) REH2-ΔN, (B) REH2-ΔNds, (C) ^H2^F1-ΔN, and (D) ^H2^F1-ΔC. Western blots of the tag in these constructs (TAP), the cytosolic marker eEF-1 (all panels), and endogenous REH2 (panels C and D) indicate a partial enrichment of the REH2 deletion constructs in the mini-mito extract. Some mitochondrial enrichment of ^H2^F1-ΔC is apparent relative to eEF-1 but the localization of ^H2^F1-ΔC is clearly compromised compared to other constructs examined.

**S3 Figure. The OB-fold in yeast helicase Prp43p (crystal structure pdb: 2XAU_B) and *T. brucei* REH2**. (A) Sequence alignment of Prp43p, REH2, and other DExH/RHA helicases. The sequences were aligned using the CDD tool in NCBI. The residue R1979 in REH2 is not conserved in the aligned sequences but the basic amino acid at position 2023 in REH2 (K704 in Prp43p) is conserved. Secondary structure elements are indicated: *α*-helix (cylinders) and β-strand (arrows). (B) homology model of R2023 in REH2 using the OB fold in Prp43p as a template.

**S4 Figure. Multi-sequence alignment of a C-terminal segment in *T. brucei* REH2 and *Drosophila* MLE DExH/RHA RNA helicases**. The alignment was generated with Clustal Omega (48). Boxes were inserted manually to improve the match between the MLE and REH2 residues. Note that H1032, K1033 and T1034 (in red), which make U-specific contacts in MLE, are aligned with H1998, R1999 and T2000 (in red) in REH2. Predicted *α*-helix (cylinders) and β-strand (arrows) in REH2 are indicated.

**S5 Figure. Sedimentation analysis of endogenous editing proteins. (A)** 10-30% glycerol gradients of freshly-made mitochondria-enriched extract from 29:13 procyclic trypanosomes. Catalase and RECC complex were used as 11 S and 20 S markers, respectively (23). Endogenous REH2, ^H2^F1 and ^H2^F2, GAP1 (GRBC2), GAP2 (GRBC1), and A2 (MP42) were examined in western blots. All panels in this figure derived from the same extract fractions. The data shown is representative of at least two panels for each protein in biological replicate gradients.

**S6 Figure. Location of the zinc-finger substitutions examined in this study. (A)** Full ^H2^F1 amino acid sequence including the location of eight C2H2 zinc-finger motifs (Znf1-to-8 highlighted in different colors)**. (B)** Zinc-finger motifs starting with the N terminal finger at the top, and the amino acid positions spanning each finger in panel A. The R/K>A substitutions in each finger that were examined in this study are marked by a dot.

**S7 Figure. RNase-resistant co-purification of tagged-^H2^F1 and endogenous ^H2^F1.** Western blots of IgG pulldowns from extracts with or without an RNaseA/T1 mix. All panels in this figure derive from the same blot. The upper blot with the tagged-^H2^F1 bait was cut below the 75 kDa marker. The middle and lower panels were divided between the 50 kDa and 37 kDa marker. The 34.4 kDa RGG2, a typical subunit of the REMC module in the RESC complex. As expected, the RNA-mediated association of RGG2 decreased with the RNase treatment.

**S8 Figure. Quantitation of steady-state RNA transcripts in the input mitochondrial extracts used in the IgG pulldowns.** Independent biological replicates (two independent cultures used in Fig. 5) with Cq average values and one standard deviation (+/-1SD, *n*=2) were plotted. dCq of steady-state RNA transcripts in lysates relative to background 18s rRNA used as reference [dCq = 2^(target Cq − ref Cq)^]. Shorter bars indicate a smaller differential versus 18s rRNA in the sample. For example, ND7 is relatively abundant compared to other transcripts in the sample (i.e., it has a lower Cq). The WT construct is induced or not (+/-). All mutants are induced. All end-point amplicons were examined in gels to confirm that they were single products during linear amplification. Abbreviation of the RNA names is as in Fig. 5B.

**S1 Table. DNA oligonucleotides, gBlocks (IDT) and synthetic RNA.**

## References

1. Cruz-Reyes J, Mooers BHM, Doharey PK, Meehan J, Gulati S. Dynamic RNA holoeditosomes with subcomplex variants: Insights into the control of trypanosome editing. WIREs RNA. 2018;e1502.

2. Koslowsky DJ, Sun Y, Hindenach J, Theisen T, Lucas J. The insect-phase gRNA transcriptome in Trypanosoma brucei. Nucleic acids research. 2013.

3. Madina BR, Kumar V, Mooers BH, Cruz-Reyes J. Native Variants of the MRB1 Complex Exhibit Specialized Functions in Kinetoplastid RNA Editing. PloS one. 2015;10(4):e0123441.

4. Kumar V, Madina BR, Gulati S, Vashisht AA, Kanyumbu C, Pieters B, et al. REH2C Helicase and GRBC Subcomplexes May Base Pair through mRNA and Small Guide RNA in Kinetoplastid Editosomes. J Biol Chem. 2016;291(11):5753–64.

5. Madina BR, Kumar V, Metz R, Mooers BH, Bundschuh R, Cruz-Reyes J. Native mitochondrial RNA-binding complexes in kinetoplastid RNA editing differ in guide RNA composition. RNA. 2014;20(7):1142–52.

6. Aphasizheva I, Zhang L, Wang X, Kaake RM, Huang L, Monti S, et al. RNA binding and core complexes constitute the U-insertion/deletion editosome. Mol Cell Biol. 2014;34(23):4329–42.

7. Huang Z, Faktorova D, Krizova A, Kafkova L, Read LK, Lukes J, et al. Integrity of the core mitochondrial RNA-binding complex 1 is vital for trypanosome RNA editing. RNA. 2015.

8. McAdams NM, Simpson RM, Chen R, Sun Y, Read LK. MRB 7260 is essential for productive protein-RNA interactions within the RNA editing substrate binding complex during trypanosome RNA editing. RNA. 2018;24(4):540–56.

9. Simpson RM, Bruno AE, Bard JE, Buck MJ, Read LK. High-throughput sequencing of partially edited trypanosome mRNAs reveals barriers to editing progression and evidence for alternative editing. RNA. 2016;22(5):677–95.

10. Simpson RM, Bruno AE, Chen R, Lott K, Tylec BL, Bard JE, et al. Trypanosome RNA Editing Mediator Complex proteins have distinct functions in gRNA utilization. Nucleic acids research. 2017;45(13):7965–83.

11. Shaw PL, McAdams NM, Hast MA, Ammerman ML, Read LK, Schumacher MA. Structures of the T. brucei kRNA editing factor MRB1590 reveal unique RNA-binding pore motif contained within an ABC-ATPase fold. Nucleic acids research. 2015;43(14):7096–109.

12. Missel A, Souza AE, Norskau G, Goringer HU. Disruption of a gene encoding a novel mitochondrial DEAD-box protein in Trypanosoma brucei affects edited mRNAs. Mol Cell Biol. 1997;17(9):4895–903.

13. Li F, Herrera J, Zhou S, Maslov DA, Simpson L. Trypanosome REH1 is an RNA helicase involved with the 3’-5’ polarity of multiple gRNA-guided uridine insertion/deletion RNA editing. Proc Natl Acad Sci U S A. 2011;108(9):3542–7.

14. Cruz-Reyes J, Mooers BH, Abu-Adas Z, Kumar V, Gulati S. DEAH-RHA helicase•Znf cofactor systems in kinetoplastid RNA editing and evolutionarily distant RNA processes. RNA Dis. 2016;3(2):e1336.

15. Jarmoskaite I, Russell R. RNA helicase proteins as chaperones and remodelers. Annu Rev Biochem. 2014;83:697–725.

16. Walbott H, Mouffok S, Capeyrou R, Lebaron S, Humbert O, van Tilbeurgh H, et al. Prp43p contains a processive helicase structural architecture with a specific regulatory domain. EMBO J. 2010;29(13):2194–204.

17. Robert-Paganin J, Rety S, Leulliot N. Regulation of DEAH/RHA helicases by G-patch proteins. Biomed Res Int. 2015;2015:931857.

18. Prabu JR, Muller M, Thomae AW, Schussler S, Bonneau F, Becker PB, et al. Structure of the RNA Helicase MLE Reveals the Molecular Mechanisms for Uridine Specificity and RNA-ATP Coupling. Mol Cell. 2015;60(3):487–99.

19. Dhote V, Sweeney TR, Kim N, Hellen CU, Pestova TV. Roles of individual domains in the function of DHX29, an essential factor required for translation of structured mammalian mRNAs. Proc Natl Acad Sci U S A. 2012;109(46):E3150–9.

20. Kelley LA, Mezulis S, Yates CM, Wass MN, Sternberg MJ. The Phyre2 web portal for protein modeling, prediction and analysis. Nat Protoc. 2015;10(6):845–58.

21. Cruz-Reyes J. MBHM, Kumar K., Doharey P.K., Meehan J., Chaparro L. Control Mechanisms of the Holo-Editosome in Trypanosomes. In: J C-R, M G, editors. RNA Metabolism in Mitochondria. Nucleic Acids and Molecular Biology. 34: Springer, Cham; 2018.

22. Jankowsky E. RNA helicases at work: binding and rearranging. Trends in biochemical sciences. 2011;36(1):19–29.

23. Hernandez A, Madina BR, Ro K, Wohlschlegel JA, Willard B, Kinter MT, et al. REH2 RNA helicase in kinetoplastid mitochondria: ribonucleoprotein complexes and essential motifs for unwinding and guide RNA (gRNA) binding. J Biol Chem. 2010;285(2):1220–8.

24. Segal DJ, Crotty JW, Bhakta MS, Barbas CF, 3rd, Horton NC. Structure of Aart, a designed six-finger zinc finger peptide, bound to DNA. J Mol Biol. 2006;363(2):405–21.

25. Chen S, Xu Y, Zhang K, Wang X, Sun J, Gao G, et al. Structure of N-terminal domain of ZAP indicates how a zinc-finger protein recognizes complex RNA. Nat Struct Mol Biol. 2012;19(4):430–5.

26. Klug A. The discovery of zinc fingers and their development for practical applications in gene regulation and genome manipulation. Q Rev Biophys. 2010;43(1):1–21.

27. Seiwert SD, Heidmann S, Stuart K. Direct visualization of uridylate deletion in vitro suggests a mechanism for kinetoplastid RNA editing. Cell. 1996;84(6):831–41.

28. Rusche LN, Cruz-Reyes J, Piller KJ, Sollner-Webb B. Purification of a functional enzymatic editing complex from Trypanosoma brucei mitochondria. EMBO J. 1997;16(13):4069–81.

29. Izzo A, Regnard C, Morales V, Kremmer E, Becker PB. Structure-function analysis of the RNA helicase maleless. Nucleic acids research. 2008;36(3):950–62.

30. Masliah G, Barraud P, Allain FH. RNA recognition by double-stranded RNA binding domains: a matter of shape and sequence. Cell Mol Life Sci. 2013;70(11):1875–95.

31. Christian H, Hofele RV, Urlaub H, Ficner R. Insights into the activation of the helicase Prp43 by biochemical studies and structural mass spectrometry. Nucleic acids research. 2014;42(2):1162–79.

32. Ye P, Liu S, Zhu Y, Chen G, Gao G. DEXH-Box protein DHX30 is required for optimal function of the zinc-finger antiviral protein. Protein & cell. 2010;1(10):956–64.

33. Guo X, Carroll JW, Macdonald MR, Goff SP, Gao G. The zinc finger antiviral protein directly binds to specific viral mRNAs through the CCCH zinc finger motifs. J Virol. 2004;78(23):12781–7.

34. Kruse E, Voigt C, Leeder WM, Goringer HU. RNA helicases involved in U-insertion/deletion-type RNA editing. Biochimica et biophysica acta. 2013;1829(8):835–41.

35. Carnes J, Soares CZ, Wickham C, Stuart K. Endonuclease associations with three distinct editosomes in Trypanosoma brucei. J Biol Chem. 2011;286(22):19320–30.

36. McDermott SM, Luo J, Carnes J, Ranish JA, Stuart K. The Architecture of Trypanosoma brucei editosomes. Proc Natl Acad Sci U S A. 2016;113(42):E6476–E85.

37. Carnes J, McDermott S, Anupama A, Oliver BG, Sather DN, Stuart K. In vivo cleavage specificity of Trypanosoma brucei editosome endonucleases. Nucleic acids research. 2017;45(8):4667–86.

38. Ammerman ML, Hashimi H, Novotna L, Cicova Z, McEvoy SM, Lukes J, et al. MRB3010 is a core component of the MRB1 complex that facilitates an early step of the kinetoplastid RNA editing process. RNA. 2011;17(5):865–77.

39. Wirtz E, Leal S, Ochatt C, Cross GA. A tightly regulated inducible expression system for conditional gene knock-outs and dominant-negative genetics in Trypanosoma brucei. Molecular and biochemical parasitology. 1999;99(1):89–101.

40. Hernandez A, Panigrahi A, Cifuentes-Rojas C, Sacharidou A, Stuart K, Cruz-Reyes J. Determinants for association and guide RNA-directed endonuclease cleavage by purified RNA editing complexes from Trypanosoma brucei. J Mol Biol. 2008;381(1):35–48.

41. Panigrahi AK, Schnaufer A, Ernst NL, Wang B, Carmean N, Salavati R, et al. Identification of novel components of Trypanosoma brucei editosomes. RNA. 2003;9(4):484–92.

42. Ammerman ML, Downey KM, Hashimi H, Fisk JC, Tomasello DL, Faktorova D, et al. Architecture of the trypanosome RNA editing accessory complex, MRB1. Nucleic acids research. 2012;40(12):5637–50.

43. Weng J, Aphasizheva I, Etheridge RD, Huang L, Wang X, Falick AM, et al. Guide RNA-binding complex from mitochondria of trypanosomatids. Mol Cell. 2008;32(2):198–209.

44. Sabatini R, Hajduk SL. RNA ligase and its involvement in guide RNA/mRNA chimera formation. Evidence for a cleavage-ligation mechanism of Trypanosoma brucei mRNA editing. J Biol Chem. 1995;270(13):7233–40.

45. Hashimi H, Zikova A, Panigrahi AK, Stuart KD, Lukes J. TbRGG1, an essential protein involved in kinetoplastid RNA metabolism that is associated with a novel multiprotein complex. RNA. 2008;14(5):970–80.

46. Harris ME, Hajduk SL. Kinetoplastid RNA editing: in vitro formation of cytochrome b gRNA-mRNA chimeras from synthetic substrate RNAs. Cell. 1992;68(6):1091–9.

47. Carnes J, Stuart KD. Uridine insertion/deletion editing activities. Methods in enzymology. 2007;424:25–54.

48. Sievers F, Higgins DG. Clustal Omega, accurate alignment of very large numbers of sequences. Methods Mol Biol. 2014;1079:105–16.

49. Marchler-Bauer A, Derbyshire MK, Gonzales NR, Lu S, Chitsaz F, Geer LY, et al. CDD: NCBI’s conserved domain database. Nucleic acids research. 2015;43(Database issue):D222–6.

