## Supplemental Figures 1 to 8 for "Protein features for assembly of the RNA editing helicase 2 subcomplex (REH2C) in Trypanosome holo-editosomes"

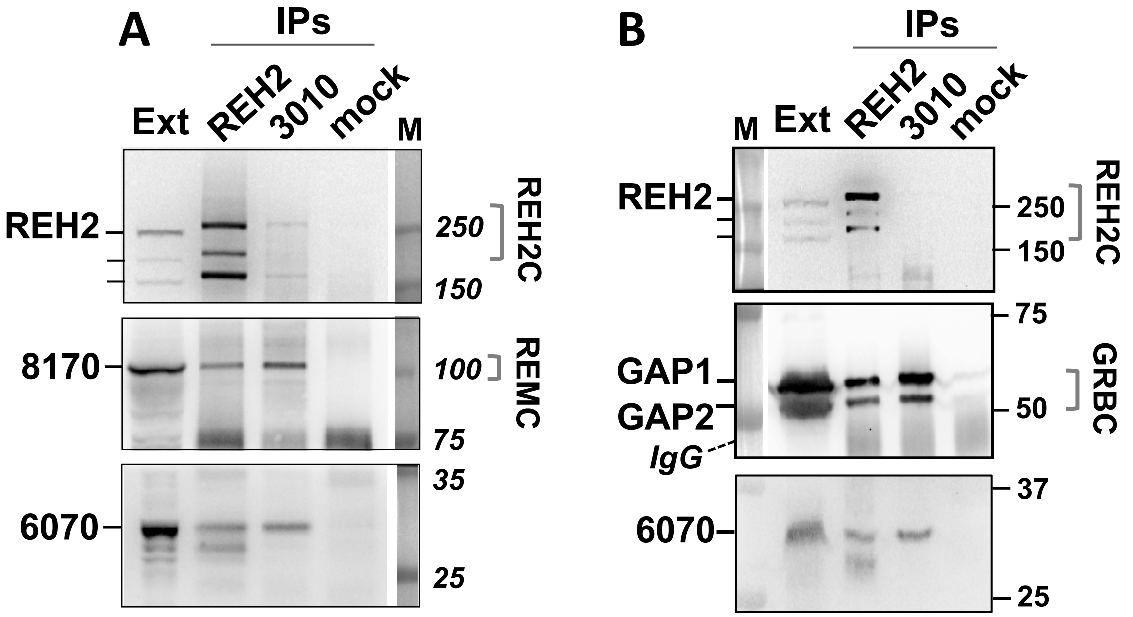
A
IPs
Ext
REH2
3010
mock
M
REH2
250
150
REH2C
8170
100
REMC
75
6070
35
25
B
IPs
M
Ext
REH2
3010
mock
REH2
250
150
REH2C
75
GAP1
GAP2
IgG
50
GRBC
6070
37
25

S1 Fig.

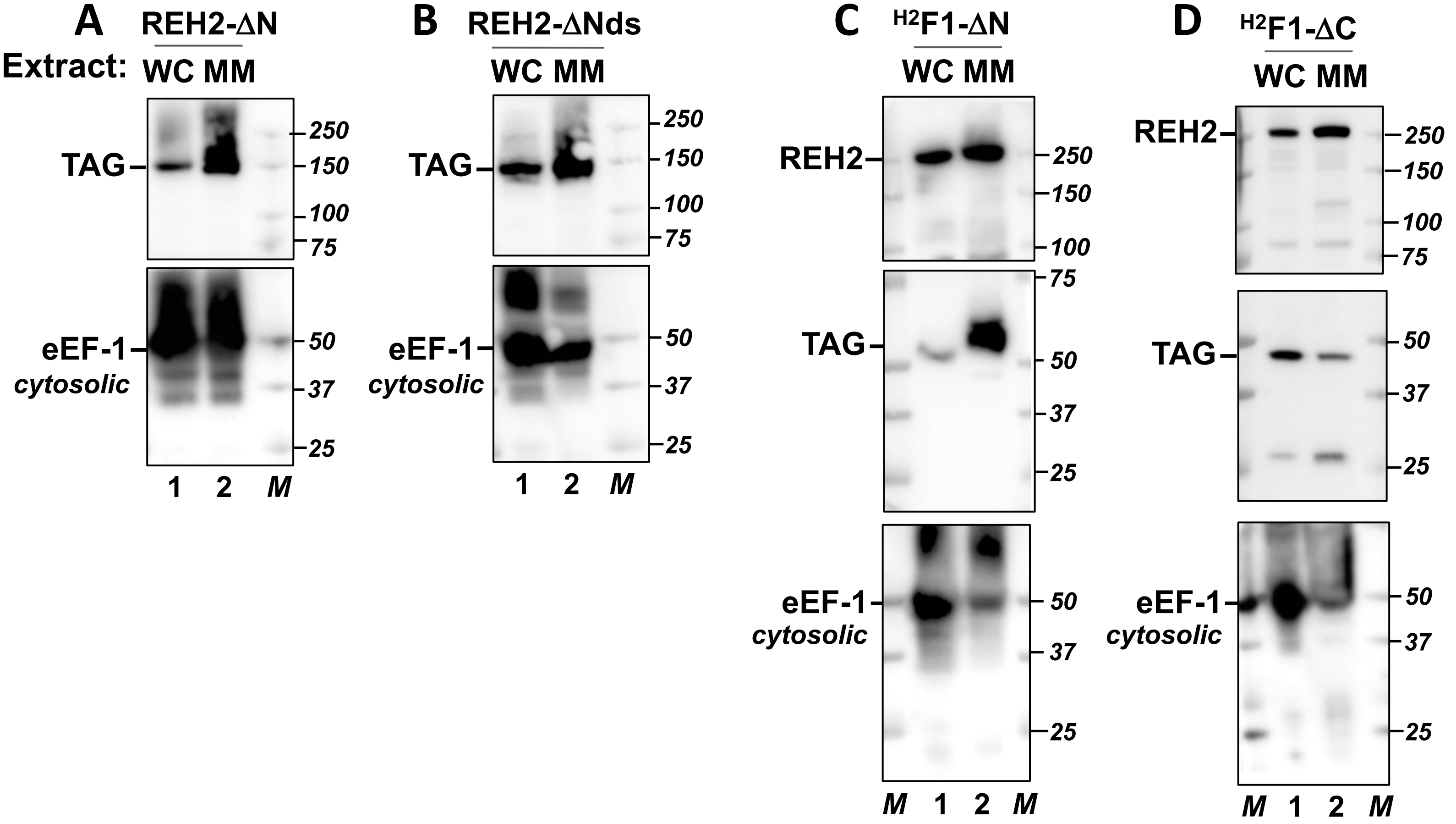
A
REH2-ΔN
Extract:
WC MM
TAG
eEF-1
cytosolic
250
150
100
75
50
37
25
1 2 M
B
REH2-ΔNds
WC MM
TAG
eEF-1
cytosolic
1 2 M
C
H2F1-ΔN
WC MM
REH2
TAG
eEF-1
cytosolic
M 1 2 M
D
H2F1-ΔC
WC MM
REH2
TAG
eEF-1
cytosolic
M 1 2 M

S2 Fig

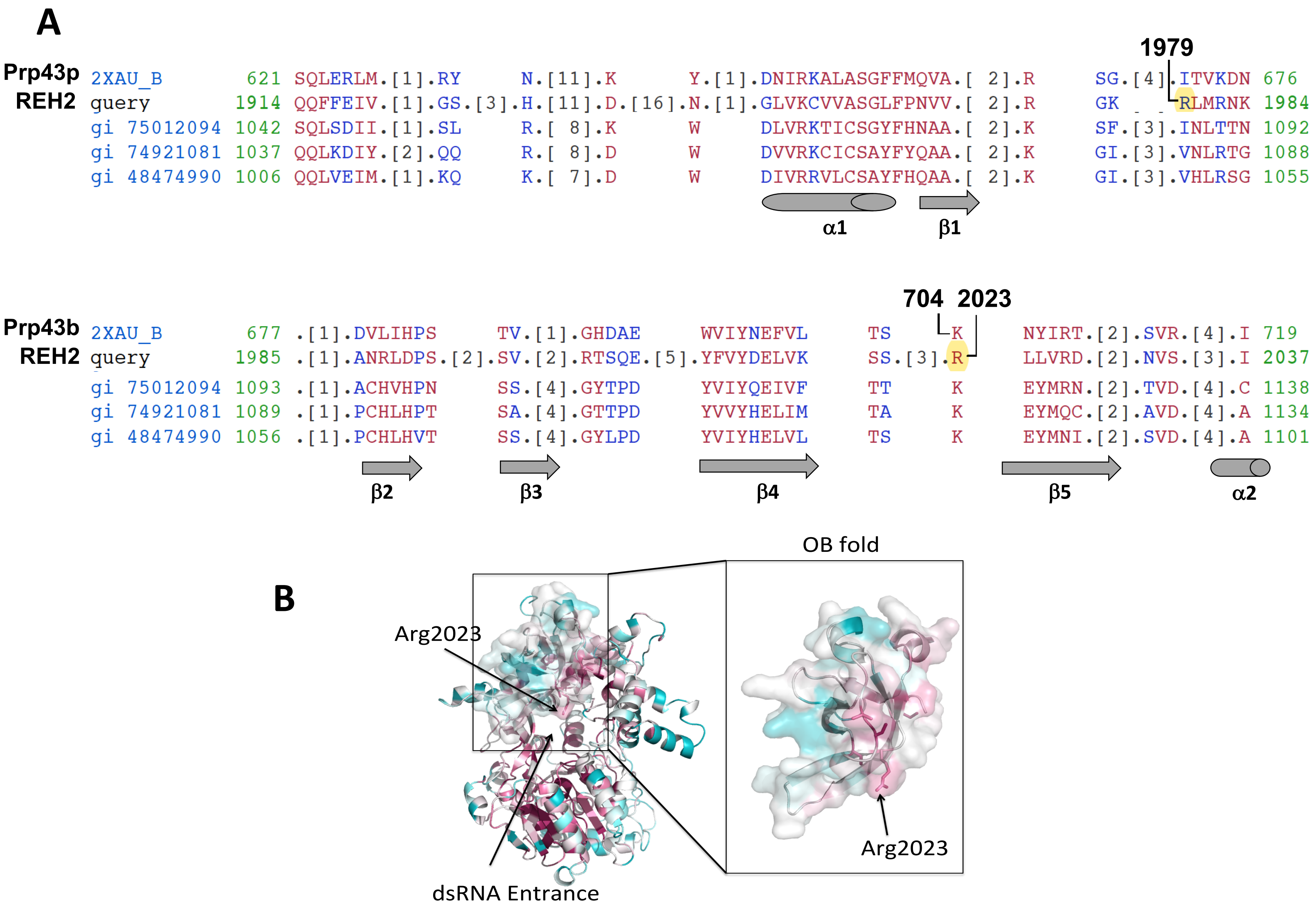
A
1979
Prp43p 2XAU_B 621 SQLERLM.[1].RY N.[11].K Y.[1].DNIRKALASGFFMQVA.[ 2].R SG.[4].ITVKDN 676
REH2 query 1914 QQFFEIV.[1].GS.[3].H.[11].D.[16].N.[1].GLVKCVVASGLFPNVV.[ 2].R GK RLMRNK 1984
gi 75012094 1042 SQLSDII.[1].SL R.[ 8].K W DLVRKTICSGYFHNAA.[ 2].K SF.[3].INLTTN 1092
gi 74921081 1037 QQLKDIY.[2].QQ R.[ 8].D W DVVRKCICSAYFYQAA.[ 2].K GI.[3].VNLRTG 1088
gi 48474990 1006 QQLVEIM.[1].KQ K.[ 7].D W DIVRRVLCSAYFHQAA.[ 2].K GI.[3].VHLRSG 1055
α1
β1
704 2023
Prp43b 2XAU_B 677 .[1].DVLIHPS TV.[1].GHDAE WVIYNEFVL TS K NYIRT.[2].SVR.[4].I 719
REH2 query 1985 .[1].ANRLDPS.[2].SV.[2].RTSQE.[5].YFVYDELVK SS.[3].R LLVRD.[2].NVS.[3].I 2037
gi 75012094 1093 .[1].ACHVHPN SS.[4].GYTPD YVIYQEIVF TT K EYMRN.[2].TVD.[4].C 1138
gi 74921081 1089 .[1].PCHLHPT SA.[4].GTTPD YVVYHELIM TA K EYMQC.[2].AVD.[4].A 1134
gi 48474990 1056 .[1].PCHLHVT SS.[4].GYLPD YVIYHELVL TS K EYMNI.[2].SVD.[4].A 1101
β2
β3
β4
β5
α2
B
OB fold
Arg2023
dsRNA Entrance
Arg2023

S3 Fig.

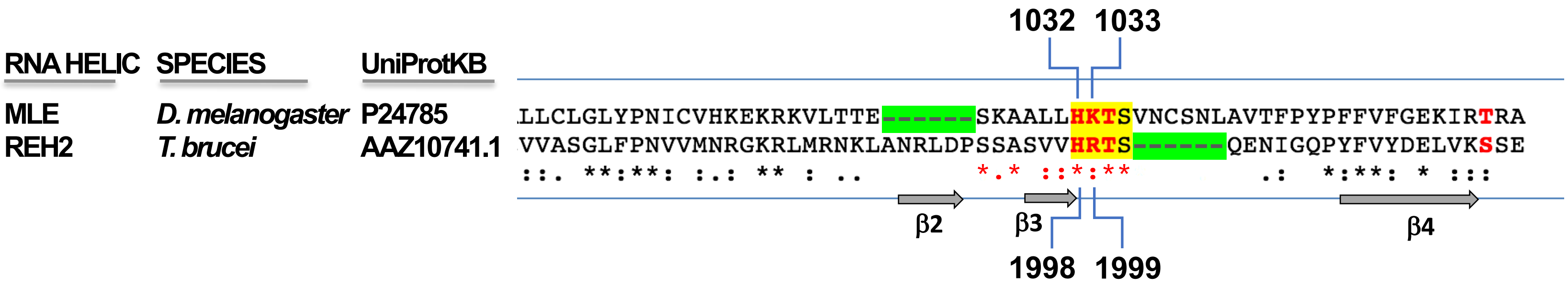
1032 1033
RNA HELIC
SPECIES
UniProtKB
MLE
D. melanogaster
P24785
LLCLGLYPNICVHKEKRKVLTTE-------SKAALLHKTSVNCSNLAVTFPYPFFVFGEKIRTRA
REH2
T. brucei
AAZ10741.1
VVASGLFPNVVMNRGKRLMRNKLANRLDPSSASVVHRTS-------QENIGQPYFVYDELVKSSE
::. **:**: :.: ** : .. *.* ::*:** .: *:**: * :::
β2
β3
β4
1998 1999

S4 Fig.

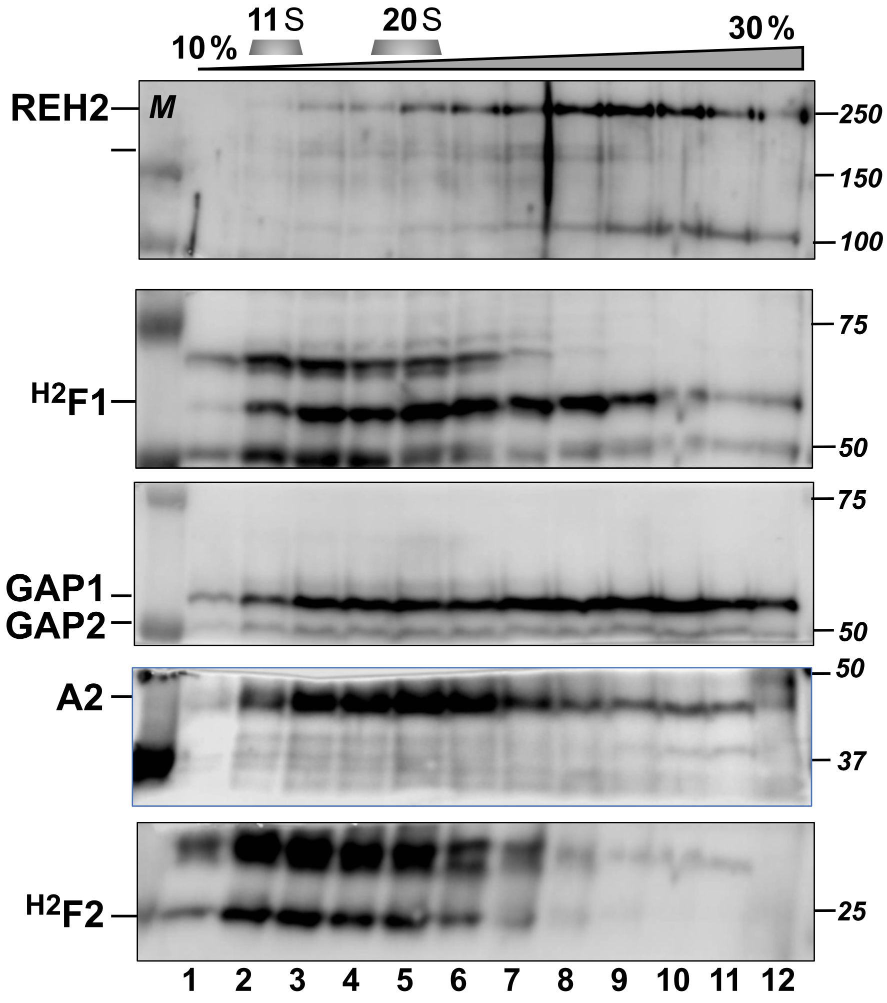
11 S
20 S
10 %
30 %
REH2
M
250
150
100
75
H2F1
50
75
GAP1
GAP2
50
50
A2
37
H2F2
25
1 2 3 4 5 6 7 8 9 10 11 12

S5 Fig.

**A**

```
1   MRRWLVASMAPQLHQLLQPVRRCHHPLRIPSVQLAAPRSHTHEDIAYASCPACSRVVHMC   Znf1
61  DMLTHLITAHRELDQTHCRKMCTERLALYERVIGVPLKKSELTSSGRRVLDFLPTVLPTG
121 YMCNWCDRRSDVYATRDKFLKHVADVHTDIDLEEVEPHVPLPPRGVVVEKSNGDGGPQPT   Znf2
181 RRLNGVVAVAEKSEPINAVPRILGISLPRGVDRPLKATAQFSDTEFPCELCNRTFNSEID   Znf3
241 LLQHLETRHPDGTAEGPAGVDSAAIADVAQFSAKEATTGGDQRVHVICDLCVSSSKVYKM   Znf4
301 PSALFSHIRFKHPNEDAAFHVERLIREQKTVSSFVCTVCQKAFASAAALDGHFNSKHAEQ   Znf5
361 GEAQNVVGRVTANNCWWCHDCEKGFSSAKGLHGHMQNKHGLSSQTHPCPACKRVFADIYS   Znf6
421 LEEHLSLQHKTIRLSDIGLLTHVKCSTCERFFLSHEDLHRHAVKHHKKDPRAPAQPFEAP   Znf7
481 TSASHVAASTSAAVPSEVEATASPQGPRKVKKRKKTTEVSEVTS                   Znf8
```

**B**

C2H2 zinc fingers

| | Highlighted sequence |
| --- | --- |
| ASCPACSRVVHMCDMLT----HLITAHRE | 48-72 |
| YMCNWCDRRSDVYATRDKFLKHVADVHTD | 120-149 |
| FPCELCNRTFNSEIDLLQ---HLETRHPD | 225-251 |
| VICDLCVSSSKVYKMPSALFSHIRFKHPN | 286-314 |
| FVCTVCQKAFASAAALDG---HFNSKHAE | 334-359 |
| WWCHDCEKGFSSAKGLHG---HMQNKHGL | 375-399 |
| HPCPACKRVFADIYSLEE---HLSLQHKT | 406-431 |
| VKCSTCERFFLSHEDLHR---HAVKHHKK | 442-468 |

R/K>A substitutions in each finger

S6 Fig.

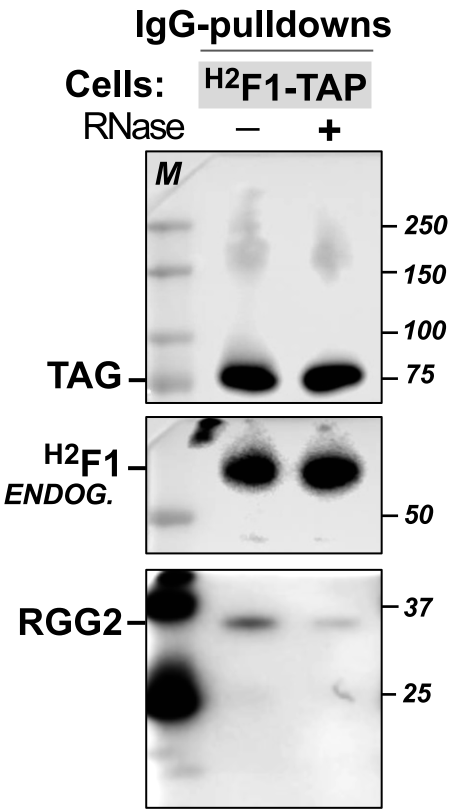
IgG-pulldowns
Cells:
H2F1-TAP
RNase
–
+
M
250
150
100
75
TAG
H2F1
ENDOG.
50
RGG2
37
25

S7 Fig.

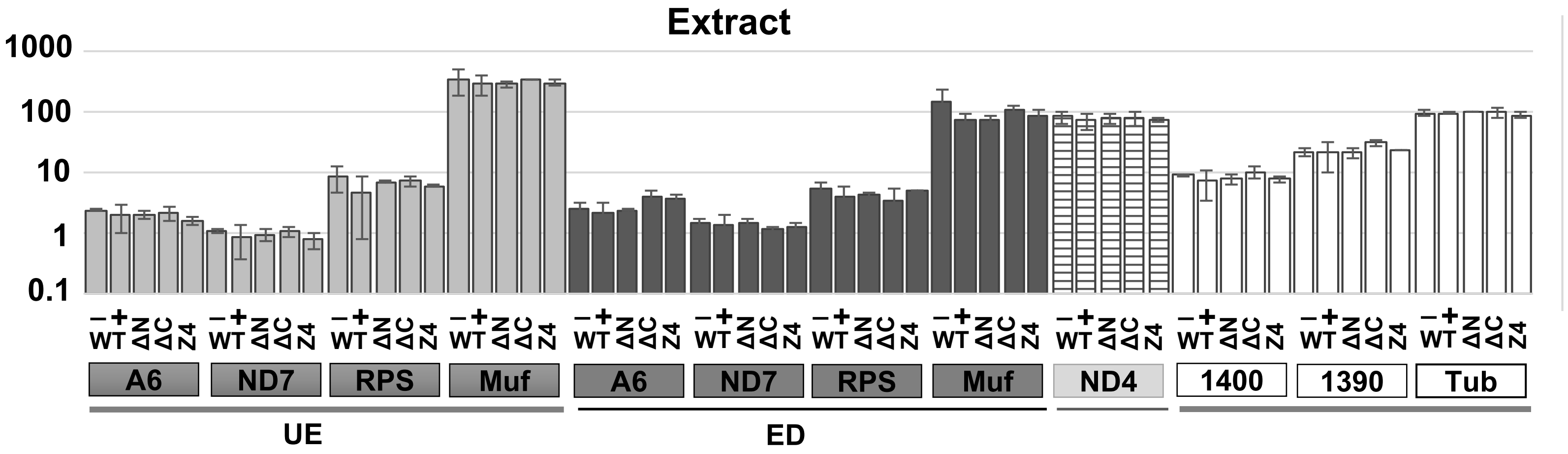
Extract
1000
100
10
1
0.1
−
+WT
ΔN
ΔC
Z4
A6
ND7
RPS
Muf
ND4
1400
1390
Tub
UE
ED

S8 Fig.
