## Supplemental Table 1 for "Protein features for assembly of the RNA editing helicase 2 subcomplex (REH2C) in Trypanosome holo-editosomes"

| REH2 mutant constructs | | | Reference |
|---|---|---|---|
| dsR (dsRBD2 K1078A/K1086A) | | | |
| *Two-step site-directed mutagenesis* | F-1045 | GGCGTAGCGTGGAAT**GCA**GAGGCCTCGCAACGC | 3 |
| | R-1046 | GCGTTGCGAGGCCTC**TGC**ATTCCACGCTACGCC | |
| K1078A primers, PCR #1 | F-1047 | GCAACGCCAGGCG**GAC**ATGCACGCC | |
| K1086A primers, PCR #2 | R-1048 | GGCGTGCAT**GTC**CGCCTGGCGTTGC | |
| ΔN (keeps dsRBD2) | | | |
| *Two fragment In-Fusion* | F-1503 | CACAAGCTTCTCGAGATGCGGGCCATACGACTAAC | |
| | R-1522 | CTGCGTTTGAAACAGTCGCACC | This study |
| | F-1523 | CTGTTTCAAACGCAGGATGCAAAAACAGTGTTGCAGCGTTAC | |
| Deletion: 989 aa (Interval: 103-3069 nt) | R-1511 | CTTTTCCATGGATCCCGAGTCTCCACCAGCCTC | |
| ΔNds (removes dsRBD2) | F-1503 | CACAAGCTTCTCGAGATGCGGGCCATACGACTAAC | |
| *Two fragment In-Fusion* | R-1522 | CTGCGTTTGAAACAGTCGCACC | This study |
| | F-1524 | CTGTTTCAAACGCAGCGCCGTGTAGAACCAGATTTCGC | |
| Deletion: 1059 aa (Interval 103-3279 nt) | R-1511 | CTTTTCCATGGATCCCGAGTCTCCACCAGCCTC | |
| ΔAOB (keeps OB) | F-1503 | CACAAGCTTCTCGAGATGCGGGCCATACGACTAAC | This study |
| Deletion: 123 aa (Interval: 6133-6501 nt) | R-1504 | CTTTTCCATGGATCCGCTGCTGGTGCCCATTAA | |
| ΔOB (removes OB) | F-1503 | CACAAGCTTCTCGAGATGCGGGCCATACGACTAAC | This study |
| Deletion: 260 aa (Interval: 5723-6501 nt) | R-1505 | CTTTTCCATGGATCCAAGGAGCTGAGGGACGC | |
| R1979A (OB) | F-1464 | GTTATGAACCGGGGGAAG**GCC**CTCATGAGG | This study |
| | R-1465 | CCTCATGAG**GGC**CTTCCCCCGGTTCATAACGACATTAGG | |
| H1998E (OB) | F-1571 | TCTGTTGTC**GAG**CGTACATCACAGGAAAATAATG | This study |
| | R-1572 | TGTACG**CTC**GACAACAGATGCCGATGATGGGTCCAG | |
| R1999E (OB) | F-1573 | TCTGTTGTCCAT**GAG**ACATCACAGGAAAATATTG | This study |
| | R-1574 | TGATGT**CTC**ATGGACAACAGATGCCGATGATGGGTCCAG | |
| H1998E/R1999E (OB) | F-1602 | TCTGTTGTC**GAGGAG**ACATCACAGGAAAATATTG | This study |
| | R-1603 | CTGTGATGT**CTCCTC**GACAACAGATGCCGATG | |
| R2023A (OB) | F-1470 | GGAATCCGAA**GCG**CTGCTCGTG | This study |
| | R-1469 | CACGAGCAG**CGC**TTCGGATTCC | |
| **H2F1 WT and mutant constructs** | | | |
| H2F1 WT | F-1487 | CATACATAAAGCTTATGCGGCGCTGGTTGGTGGC | This study |
| | R-1490 | ATCAGCAGGATCCCGACGTCACCTCACTTACC | |
| ***Canonical cysteine residues*** | | | |
| Z5 C-to-A | F-1582 | GTTTGCCAGGCGGCTTTCGCTTCCGCTG | This study |
| | R-1583 | AGCGAAAGCCGCCTGGCAAACCGTGC | |
| Primers PCR#1 | F-1584 | CTTTAATAGCGCGCATGCGGAACAAGG | |
| Primers PCR#2 | R-1585 | GTTCCGCATGCGTATTAAAGTGTCC | |
| ***Variable basic residues*** | | | |
| Z1 R/K-to-A | 1590 | gBlock — Sequences available upon request | This study |
| Z2 R/K-to-A | 1591 | gBlock | |
| Z3 R/K-to-A | 1592 | gBlock | |
| Z4 R/K-to-A | 1622 | gBlock | |
| Z5 R/K-to-A | 1623 | gBlock | |
| ***N- and C- truncations*** | | | |
| ΔN | F-1503 | CACAAGCTTCTCGAGATGCGGGCCATACGACTAAC | This study |
| *Two fragment In-Fusion* | R-1522 | CTGCGTTTGAAACAGTCGCACC | |
| The MLS derives from REH2 (see footnote) | F-1609 | CTGTTTCAAACGCAGGGGCCCGCAGGTGTTGAC | |
| Deletion: 255 aa (Interval: 1-765 nt) | R-1608 | CTTTTCCATGGATCCGGGCCCCTCTGCAGT | |
| ΔC | F-1607 | CACAAGCTTCTCGAGATGCGGCGCTGG | This study |
| Deletion: 267 aa (Interval: 772-1572 nt) | R-1608 | CTTTTCCATGGATCCGGGCCCCTCTGCAGT | |
| **mRNA/gRNA duplex** | | | |
| gRNA | | | |
| Template in PCR | 1354 | AAGCAGAAGAGATACGTTTAAAAAAAAAAAAAAAAATTATCATACCACTGTAAA | |
| | | ACTGATTTCGTCATTGGAGTTATAGTTATATCCTATAGTGAGTCGTA | 23 |
| Primers in PCR | F-1356 | AAGCAGAAGAGATACGTT | |
| | R-1385 | TAATACGACTCACTATAGGATATACTATAAC | |
| mRNA | 566 | GAGAGAGGAGAGAAGAAAGGGAAAGUUGUGAUUUUGGAGUUAUAGAAUACUUACCUGGCAUC | 23 |
| Synthetic fragment mA6 11-72 | | | |
| **Cloning recombinant REH2** | | | |
| | F-REH2 & | AGAAGGAGATATACCATGGCCATGTTTCAAACGCAGGAAATTAC | This study |
| Amino acids 30-2167 | R-REH2 # | GGCTTTGTTAGCAGCCGGATCCTCATTAGTGATGGTGATGGTGATGCGGCGAG | |
| | | TCTCCACCAGCCTCAGC | |

& full name: pET15bNCO1REH2fwd

### full name: pET15b2stop6HisProlineREH2Rev

MLS: mitochondrial leader sequence fragment (Primers: F-1503, R-1522)

The undeline indicates the mutated nucleotides in the indicated primers.
